# Impact of low-calorie sweeteners on gut bacteria is modulated by common xenobiotics

**DOI:** 10.1101/2025.03.28.645995

**Authors:** Sonja Blasche, Vinita Periwal, Nonantzin Beristain Covarrubias, Anna Lindell, Indra Roux, Stephan Kamrad, Rob Bradley, Rui Guan, Hilal Ozgur, Bini Ramachandran, Vladimir Benes, Kiran Raosaheb Patil

## Abstract

The gut microbiota is implicated in adverse effects associated with low-calorie sweeteners. Yet, the direct impact of sweeteners on gut bacteria remains largely uncharacterized. Here we report interactions between 25 phylogenetically diverse gut bacterial strains and 39 commercially used sweeteners. We tested these sweeteners individually and in combination with four commonly co-consumed compounds, viz., advantame, caffeine, vanillin, and duloxetine. Three quarters of the tested sweeteners individually impacted growth of at least one bacterial strain. Further, over 100 interactions were found between sweeteners and the four co-consumed compounds. Isosteviol, a commonly used sweetener, and duloxetine, an antidepressant, synergistically inhibited *Roseburia intestinalis*, a bacterium previously linked to glucose homeostasis, and *Parabacteroides merdae*, a prevalent commensal linked to healthy microbiota. Proteomic, metabolomic, and genetic analyses indicate altered small molecule transport underpinning this sweetener-drug synergy. The isosteviol-duloxetine combination also modulated metabolism of a synthetic gut bacterial community leading to increased toxicity to HeLa cells and altered secretion of inflammation modulatory cytokines IL-6 and IL-8 by Caco-2 cells. Together, our data bring forward the prevalence of interactions between low-calorie sweeteners and common xenobiotics.

## Introduction

Utilization of low-calorie and non-nutritional sweeteners was anticipated to enhance population health^1^. However, commonly used sweeteners have now been associated with type II diabetes, obesity, cardiovascular disease, increased cancer risk, and infection^2–6^. Further, studies are linking the consumption of artificial sweeteners with pathogenicity^7^, sucralose with genotoxicity^8^, and aspartame with neurotoxicity and cancer^5^. Some effects of sweeteners on human health are likely to be mediated through their interaction with the gut microbiota^9,10^. Although several studies have reported interactions between sweeteners and gut microbiota^2,4,11–22^, systematic studies testing for direct interactions between phylogenetically diverse gut bacteria and sweeteners are scarce. Further, gut bacteria are likely to encounter sweeteners in conjunction with other xenobiotics such as medications or food additives. Recent studies have highlighted the importance of studying the effects of co-consumed food compounds on gut bacteria^23^. However, knowledge of the interactions between sweeteners and xenobiotics, along with the underlying mechanisms, remains limited.

Commercial sweeteners are widely used in processed foods like soft drinks, candy, chewing gum, and cereal, and are frequently co-formulated with other sweeteners, the flavour compound vanillin, and the neurostimulator caffeine. Sweeteners also function as excipients in various pharmaceuticals^24^ such as antihypertensives and are co-ingested with long-term medications like antidepressants that are consumed by millions of people daily^25^. The exposure of a significant portion of the population to sweeteners in conjunction with other food compounds and drugs raises questions about the interactive effects on gut bacteria. To our knowledge, sweeteners and common xenobiotics have not been tested systematically for combinatory effects on human gut bacteria, which can be highly relevant as exemplified by multi-antibiotic-resistance phenotype in *E. coli* resulting from combination of antimicrobial compounds with vanillin^26^.

Previous studies on the impact of sweeteners on the gut microbiota and gut bacteria^9,22^ suggest prevalent effects across bacterial phylogeny. Supplementary table 1 summarizes a variety of prior in vivo and in vitro studies. However, these studies do not necessarily imply direct effects since emergent effects may arise in a community context as observed in the case of therapeutic drugs^27^, or from mixture effects involving co-consumed foods or compounds.

In this study, we conducted a comprehensive screening of all commercially available sweeteners and sugar substitutes, both natural and synthetic (totalling 39), to assess their effects on the growth of 25 strains of human gut bacteria (Supplementary tables 2 and 3a). The 25 bacteria were selected to cover a broad phylogenetic and metabolic diversity representative of the healthy human gut microbiota^28^, and to span commensals, probiotic species, and opportunistic pathogens. The 39 sweeteners cover all commercially used sweeteners that could be purchased as pure compounds. Additionally, we examined the combined effects of sweeteners with four commonly co-consumed compounds: advantame, caffeine, vanillin, and duloxetine. The four combination compounds were chosen due to their widespread usage. Advantame is a non-caloric artificial sweetener that is increasingly being used and is known to reach the gut^29,30^. Caffeine and vanillin are found in, among many other products, beverages, cakes, candies and in case of vanillin even in baby formula^31^. Duloxetine is a commonly used antidepressant drug^32^, which has been shown previously to have significant effects on metabolite secretion in gut microbes^33^. These compounds are commonly consumed by millions of individuals ^34,35^. This, together with the high usage of sweeteners indicates that substantial number of people are co-consuming sweeteners and the xenobiotics chosen in our study.

## Results and Discussion

### Three quarters of sweeteners negatively impact gut bacterial growth

We first investigated growth effects of 39 individual sweetener molecules on the 25 gut bacterial strains (975 sweetener-bacteria pairs in total, Fig. 1a, Methods). To ensure reliable growth of the selected phylogenetically diverse species, we chose mGAM (modified Gifu anaerobic medium broth, HyServe) as growth medium. This medium supports growth of all selected species in monocultures, and has been used in previous studies^28,36,37^ enabling comparative analysis. Furthermore, mGAM medium closely aligns with the *in vivo* relative abundance in healthy subjects compared to monoculture growth^28^.

**Fig. 1.**
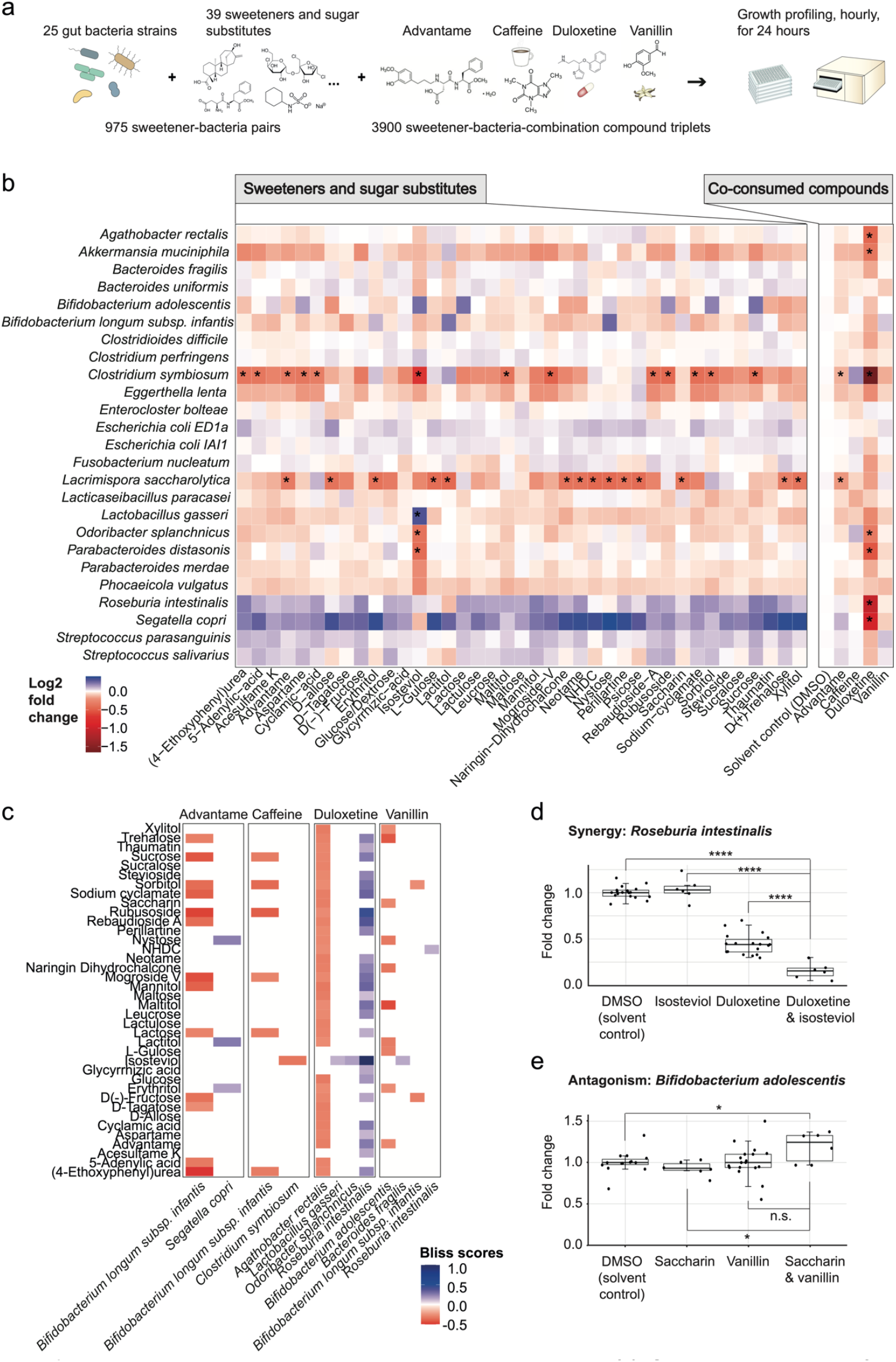
Interactions between sweeteners and gut bacteria. **(a)** Schematic overview of the sweetener-bacteria screening set-up. **(b)** Identified interactions between sweeteners and human gut bacteria. All compounds were tested at 50 µM concentration. Interactions were deemed significant (marked with *) when, a) bh-corrected^39^ p-value <0.05, b) > 20% change in growth (area under the growth curve, AUC) compared to the solvent control, and c) confirmed in an independent experiment including additional concentrations as described in the main text. **(c)** Sweeteners-xenobiotic interactions impacting gut bacterial growth. Bliss score for the significant interactions (bh-corrected p-value<0.05, and abs (log2 fold change) >0.32) are shown in the colormap. **(d)** Example of synergistic interaction between isosteviol and duloxetine affecting growth of R. intestinalis **(e)** Example of an antagonistic interaction between saccharin and vanillin affecting growth of B. adolescentis. Significance (p-val: *<0.05, **<0.01, ***<0.001, ****<0.0001) in (d) and (e) was determined using Welch’s t-test.

To enable comparison across all sweetener compounds, we used concentration of 50 µM which we estimated within the range of concentrations relevant for the colon (supplementary table 3b). Significant sweetener-bacteria interactions (p<0.05 and at least 20% change in growth) were re-tested in an independent batch at 25, 50, 100, 200 and 400 µM concentrations. Interactions significant in both cases and showing effect in at least two concentrations were considered true positives. We observe high overlap between the first and the second experiment (>90% overlap, 30 interactions) (supplementary figures 1-2, supplementary tables 4a-d, Fig 1b).

While some direct sweetener-bacteria interactions such as between stevia glycosides and lactobacilli have been described ^38^ (overview in supplementary table 1), the 30 direct interactions identified in this study have not been previously reported to our knowledge. *Clostridium symbiosum* or *Lacrimispora saccharolytica* were the most sensitive species being affected by many compounds. On the sweetener side, isosteviol was the most potent compound, inhibiting the growth of three bacteria (*C. symbiosum, Odoribacter splanchnicus*, and *Parabacteroides distasonis*) while promoting the growth of one, *Lactobacillus gasseri*.

### Antagonistic and synergistic effects of sweetener-xenobiotic combinations

To assess whether other consumed components can alter the effect of sweeteners on gut bacteria, we screened 156 sweetener-xenobiotic combinations (39 sweeteners and 4 xenobiotics) against 25 bacteria (3900 interactions in total). We also tested 12 sweetener combinations with commonly used drugs (300 interactions), reflecting typical tablet formulations. For example, acesulfame potassium is frequently combined with ibuprofen, acetaminophen, and cetirizine. Inter-compound interactions were evaluated using the Bliss model of independence^40,41^ with significance determined by a p-value threshold of 0.05 and a 20% growth change relative to each compound alone. (Methods, supplementary tables 4a-f and 5a-b, supplementary figure 3, 4).

The combination screen revealed 102 interactions involving nine gut bacteria (Fig. 1c, p-value <0.05 and log2 fold change (L2FC) <-0.32 and >0.32), 68 of which were antagonistic and 34 synergistic. To our knowledge, none of the interactions have been reported before. The higher prevalence of antagonistic interactions is consistent with those observed for drug combinations^26^, and with the expectation that the distinct compounds generally target distinct pathways. The sweetener isosteviol was notable in our results, exhibiting combinatory effects with three of the four tested xenobiotics and affecting five bacterial species. The strongest synergy was observed between isosteviol and duloxetine against *R. intestinalis* (Fig. 1d), a bacterium positively associated with health and the prevention of intestinal inflammation^42^. Combinations showing antagonistic interactions included vanillin and saccharin, which are frequently co-used in processed food products (Fig. 1e).

### Duloxetine-isosteviol combination reduces community diversity

To assess how impact on individual species can manifest at community level, we tested the effect of the duloxetine-isosteviol combination in a synthetic community assembled from the 25 gut bacteria used in the screen. The assembled community was passaged five times in the presence of DMSO (solvent control), isosteviol, duloxetine, and the isosteviol-duloxetine combination (50 µM each, Methods). Community composition was assessed directly after community assembly (p0) and after each passage (24 hours growth, p1-p5), (Fig. 2a, supplementary tables 6a, b). All assemblies exhibited early-transfer compositional dynamics that stabilized around passage p3-p4 (Fig. 2b, supplementary figure 5, supplementary table 7a), which is consistent with previous experiments with *in vitro* assembled gut bacteria communities^36,43^. A significant decrease in overall species diversity was observed for all compound-treated samples (Fig. 2-c, supplementary table 7b) and most markedly in case of the isosteviol-duloxetine combination (Fig. 2d-e). Samples challenged with the combination showed decreased relative abundance (as compared to the DMSO control) for several species, among them *R. intestinalis, P. merdae, S. copri, B. uniformis* and *P. vulgatus* (supplementary figure 6 and supplementary table 8). The relative abundance of *R. intestinalis* in the combination treated communities was significantly altered in p5, whereas that of *P. merdae* was impacted in all passages (Fig. 2f).

**Fig. 2.**
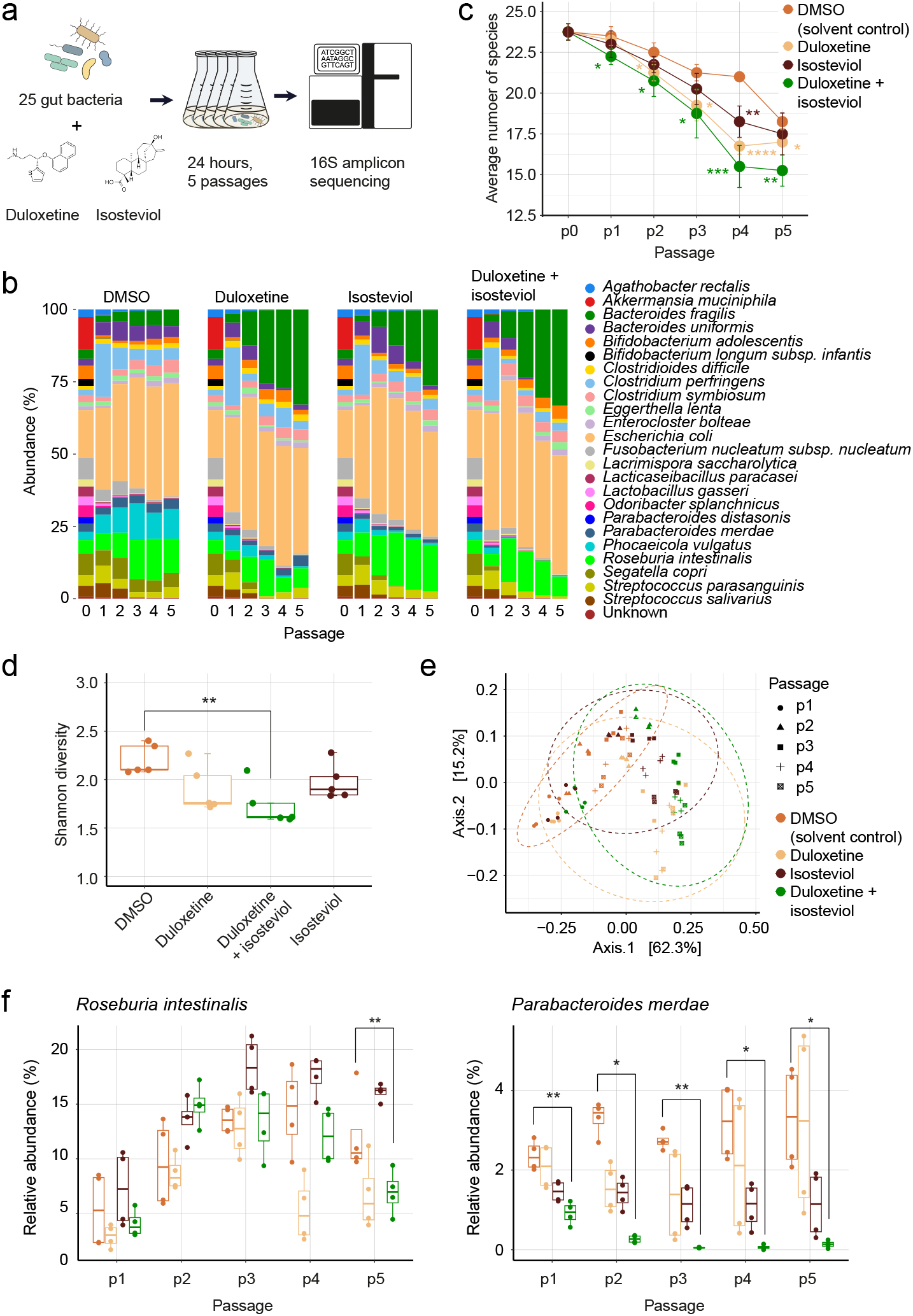
Impact of duloxetine-isosteviol combination on a synthetic gut bacterial community. **(a)** Experimental approach. **(b)** Community compositional dynamics in compound treated communities assessed using 16S rRNA analysis. **(c)** Number of surviving species during serial transfer in the control and the compound-treated communities. Lost species cutoff, <10 reads. **(d)** Changes in alpha diversity (Shannon index) between compound-treated samples. **p-val = 0.0089 (ANOVA) **(e)** Results of principle coordinate analysis showing Bray-Curtis beta-diversity across all passages and treatment groups. **(f)** Changes in the relative abundance of R. intestinalis and P. merdae during community passaging. n=4, p-val (Student’s t-test): *<0.05, **<0.01, ***<0.001, ****<0.0001.

**Fig. 3.**
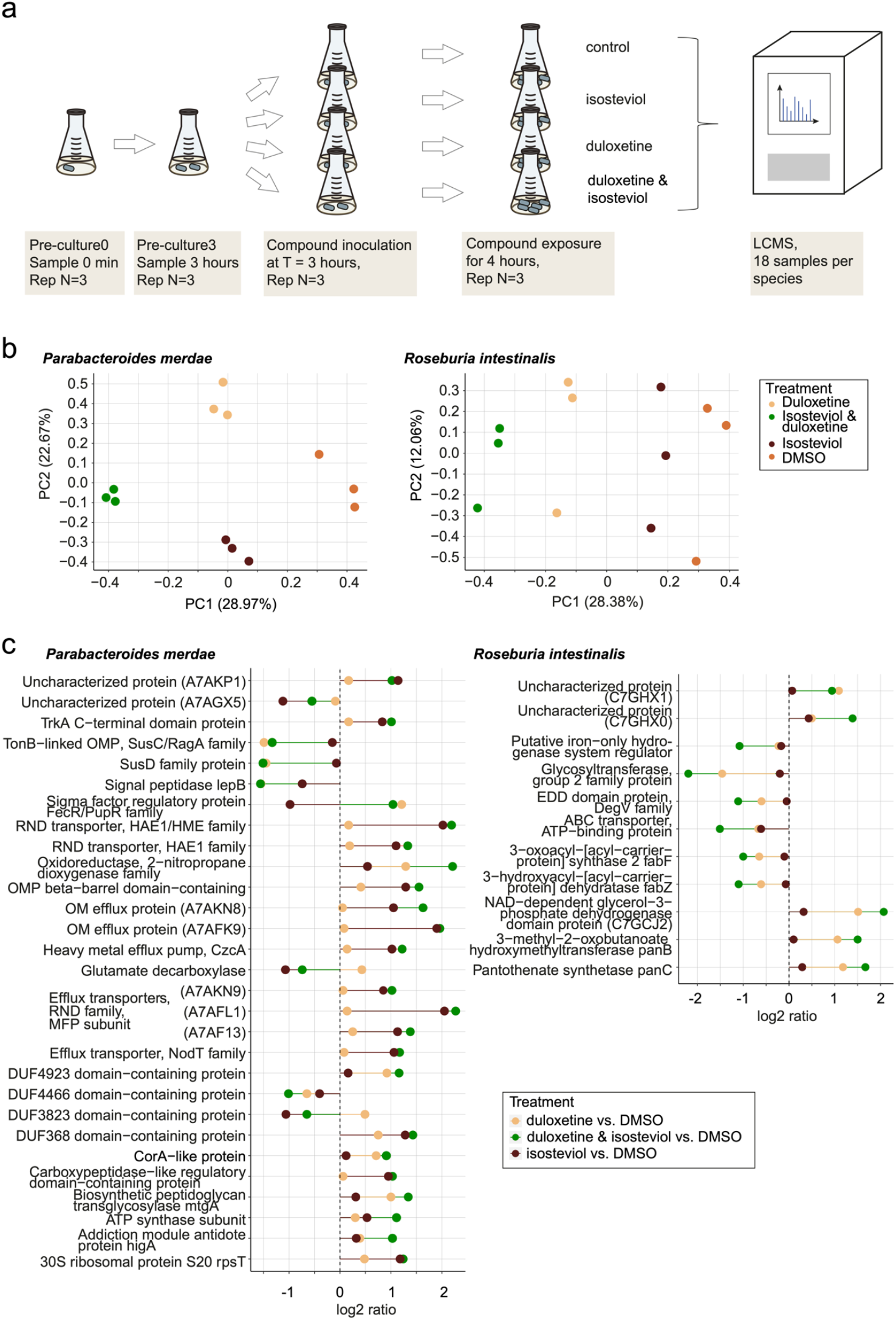
Proteomics analysis provides insights into synergistic effects of isosteviol-duloxetine combination. **(a)** Experimental design. To align cultures, bacteria were pre-grown until the logarithmic phase (OD of ∼0.6) was reached, then split into 4 aliquots and compound treatment was started and performed for 4 hours, before cultures were harvested for proteomics. **(b)** PCA showing differences between compound treated bacteria and control (DMSO). **(c)** Differently expressed proteins in treated samples normalised to DMSO-treated samples (control), N=3, cut-off: log2 FC= 1 and -1. OM, outer membrane, OMP, outer membrane protein

**Fig. 4.**
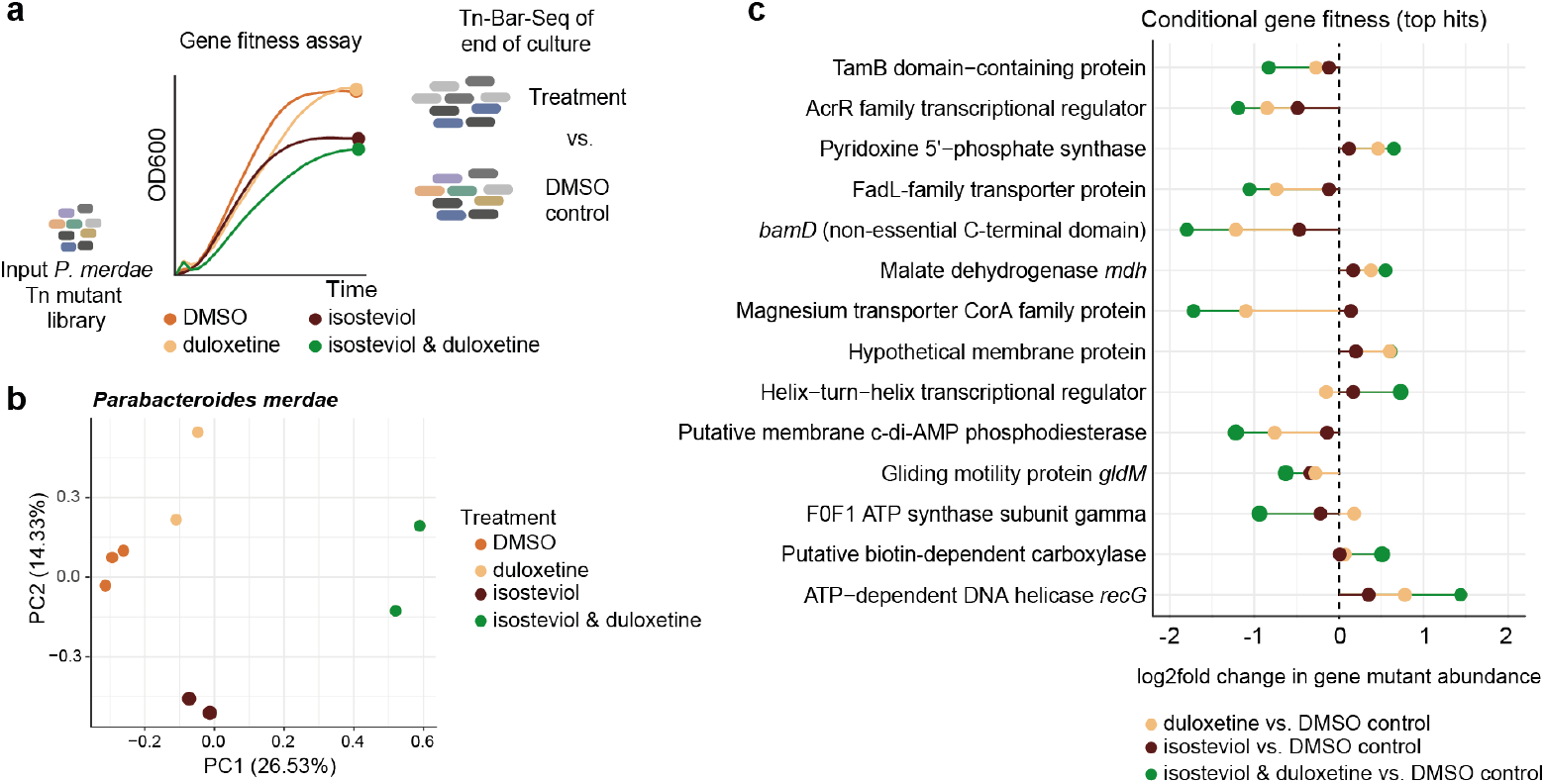
Genetic elements underlying isosteviol-duloxetine synergy. **(a)** Experimental design and representative Tn library growth curves as measured from a 100 µl sub sample. A pooled genome-wide mutant library of P. merdae was used to inoculate mGAM with the different chemical treatments or a DMSO control, following a similar growth trajectory than the WT strain. The changes in mutant abundance at end of culture were analysed by deep sequencing of the barcoded transposons (Tn-Bar-Seq, Methods). **(b)** Principal component analysis of mutant gene abundance across treatments shows a synergistic effect in the combination. **(c)** Conditional gene fitness by TnBarSeq for each condition, relative to the DMSO control. N=2 treatment, N=3 DMSO, p-adj. <0.05 and absolute log2fold >0.5 in at least one condition.

**Fig. 5.**
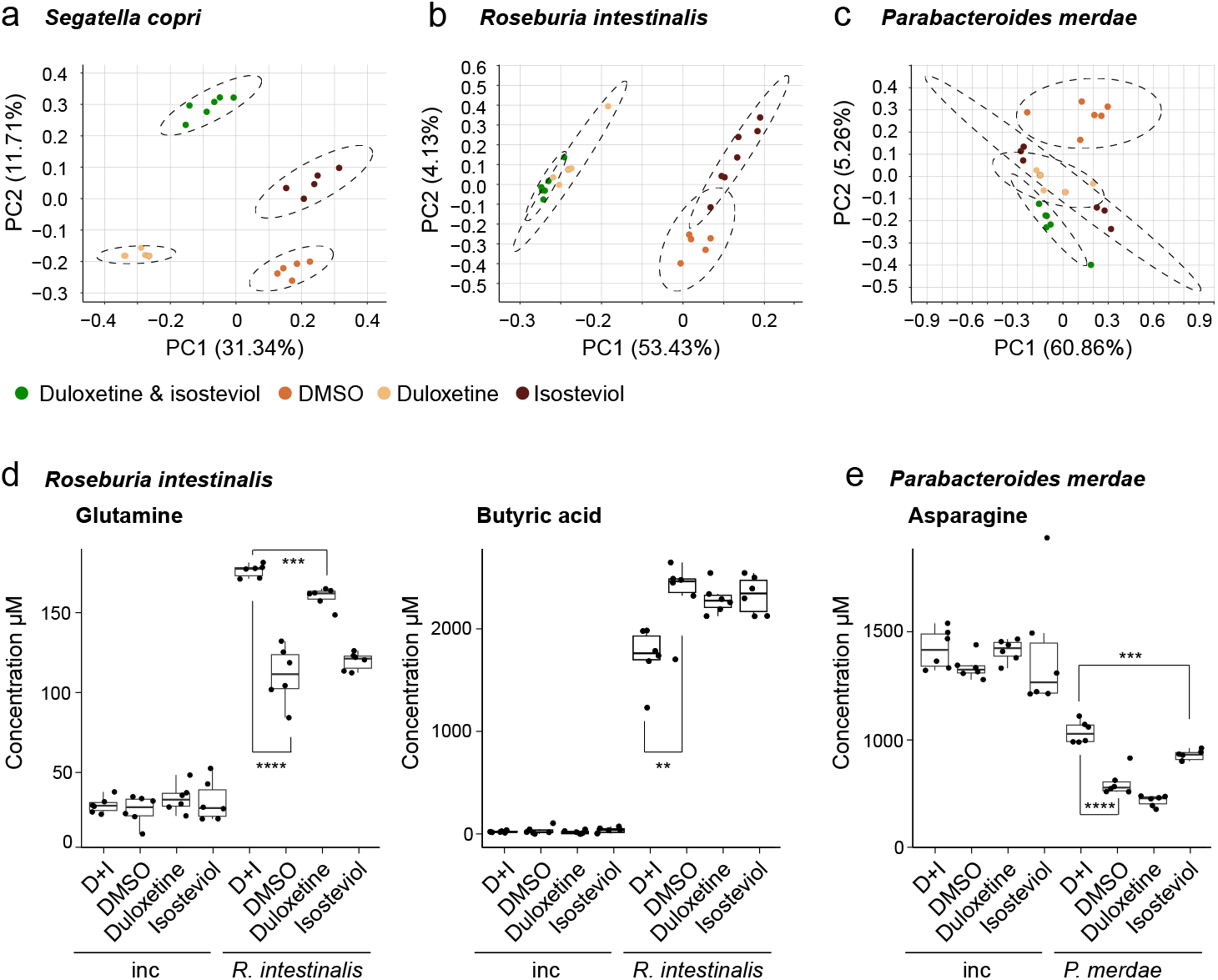
Isosteviol and duloxetine alter metabolic physiology. Altered metabolite production in culture supernatant of (a) S. copri, (b) R. intestinalis and (c) P. merdae upon exposure to isosteviol, duloxetine and a combination of both. Metabolites (captured in negative mode, using LC-MS, qTOF) cluster in a treatment-dependant way. Biological replicates: N=6, DMSO= mock, solvent control. (d) Changes of primary metabolites glutamine, butyric acid in R. intestinalis and (e) asparagine in P. merdae upon 24 hours co-exposure to duloxetine and isosteviol. D+I, duloxetine + isosteviol; inc, incubation control (medium plus compounds). Significance was determined using student’s t-test (*<0.05, **<0.01, ***<0.001, ****<0.0001).

**Fig. 6.**
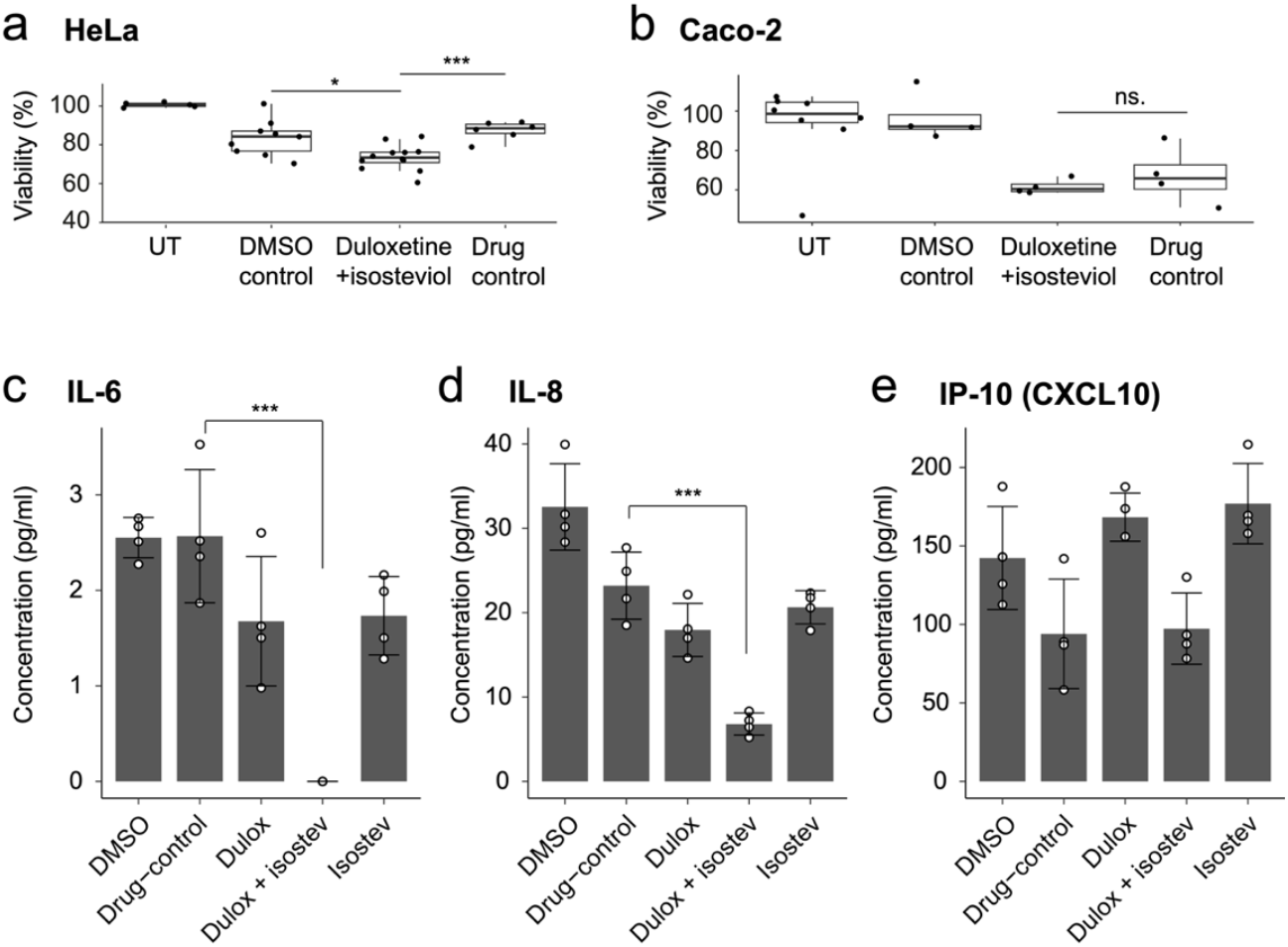
Metabolic response of gut bacterial community to isosteviol-duloxetine combination impacts cytotoxicity and cytokine secretion. (a) Spent medium of duloxetine and isosteviol co-treated communities show increased toxicity in HeLa cells, but not in (b) Caco-2 cells. And secretion of the cytokines (c) interleukin-6 (IL-6), (d) interleukin-8 (IL-8) and (e) the interferon-γ inducible protein 10 kDa (CXCL10 or IP-10) by Caco-2 cells as a response to contact with spent medium. IL-6 and IL-8 secretion is significantly reduced for the supernatant of duloxetine and isosteviol co-treated communities, as compared to the DMSO and drug control. The 25-bacteria community was treated with duloxetine (dulox), isosteviol (isostev), and duloxetine + isosteviol. Controls are DMSO (solvent, 0.2%) treated community and the supernatant of the solvent-treated community with 50 µM duloxetine and isosteviol, added after harvesting (drug control). N>6, significance was determined by Welch’s t-test. *pvalue <0.05, ***pvalue <0.001, UT untreated.

In the case of *P. merdae*, its relative abundance in the combination-treated community was significantly lower in many passages compared to treatments with either compound alone (supplementary table 9), indicating a strong combination effect. Correspondingly, ANOVA revealed a significant change in *P. merdae* abundance, while no such effect was observed for R. intestinalis (supplementary table 10). Notably, *R. intestinalis* exhibited a synergistic effect in monoculture, but not in community, whereas the reverse was true for *P. merdae*. This contrast suggests emergent effects within the community that extend beyond the direct impacts of isosteviol and duloxetine on isolated gut bacteria. Additionally, of the 25 bacteria screened, 24 showed reduced growth in monoculture after treatment with duloxetine and isosteviol, while 7 benefited in community settings. These community-effects may involve altered metabolic activity of community members as previously observed with human-targeted drugs^33^.

### Proteomic changes underlying synergy between isosteviol and duloxetine

To gain molecular insights into the isosteviol-duloxetine interaction, we assessed changes in protein abundances in *R. intestinalis* and *P. merdae* in response to the combination and individual compounds. *R. intestinalis* showed a strong synergistic effect in monoculture, while *P. merdae* exhibited an additive effect in monoculture (supplementary figure 7) and an interaction effect in the community. To ensure that the bacteria were at the same growth phase when treated, we pre-grew each species for ∼3 hours to allow reaching the logarithmic growth phase and performed proteomic analysis following 4 hours exposure (Fig. 3a). This experimental design allowed reducing growth-based differences that can be more pronounced than those resulting from exposure to different compounds^44^ (supplementary figure 8).

Principal component analysis of proteomic changes showed good concordance with the observed synergistic growth effects (Fig. 3b). Overall, the protein abundance changes in the combination-treated cells differed from the null expectation of a purely additive effect of single compound treatments (supplementary figure 9). Eleven proteins in *R. intestinalis* and 29 in *P. merdae* exhibited differential abundance upon co-exposure (Fig. 3c; supplementary tables 11-14). There was no overlap between the responding proteins from the two species suggesting distinct underlying mechanisms. The response of *P. merdae* involved 16 cell envelope associated proteins, while *R. intestinalis* exhibited many changes in the vitamin B5 pathway, i.e. increased abundance of PanB, PanC and C7GCJ2 - a putative PanG based on gene location and similarity in the pantothenate operon^45^ - and fatty acid metabolism (Fig. 3c).

To test whether vitamin B5 metabolism could modulate the synergy between isosteviol and duloxetine, we tested 20 different metabolic supplements and mixes for both *R. intestinalis* and *P. merdae*. The tested supplements included calcium pantothenate, naftidrofuryl-oxalate (an inhibitor of vitamin B5 synthesis), a mix of essential vitamins and minerals, a mixture of amino acids, sodium chloride, proteinase K, beta-alanine, butyrate, succinate, acetate, coenzyme A, ATP, additional glucose, folate, trimethoprim (an inhibitor of folate biosynthesis), all in different concentrations and combinations. However, none of these factors influenced the sensitivity of *R. intestinalis* or *P. merdae* to isosteviol, duloxetine, or their combination (see supplementary figure 10). Vitamin B pathway proteins are thus not likely to be the primary targets of duloxetine or isosteviol.

In the case of *P. merdae*, we found 29 proteins uniquely responding to the compound combination, albeit with an imbalance between the up- (21) and the down- (5) regulated proteins (Fig. 3c). Three were upregulated by one compound and downregulated by the other (sigma factor regulatory protein FecR/PupR family, glutamate decarboxylase and DUF3823 domain−containing protein). Among the up-regulated proteins, twelve are subunits of cell envelope proteins, transporters, or efflux pumps. Among the five down-regulated proteins are the SusC and SusD subunits of the saccharide uptake system, a transporter for the uptake of large nutrients through the outer membrane^46^. CorA-like protein, which is a putative magnesium transporter, had increased abundance, albeit just below the two-fold cut-off, upon combined duloxetine-isosteviol exposure. Collectively, these changes suggest *P. merdae* uses metabolite transport to counter the inhibitory combination of isosteviol and duloxetine.

### A forward genetic screen reinforces the role of transporters in sweetener toxicity

To identify genetic determinants underlying bacterial response to single and co-compound treatment, we used an in-house barcoded transposon mutant library of *P. merdae* (Tn-Bar-Seq) spanning all non-essential genes (circa 3000) (Voogdt et al., unpublished). The pooled library includes over 50 unique barcoded insertion strains per gene, which are quantified using barcode next-generation sequencing to assess gene fitness changes across perturbations (Methods).

The *P. merdae* mutant library was grown under isosteviol, duloxetine or co-exposure and the resulting compositional frequency was compared to that treated with the vehicle (DMSO) (Fig. 4a). In line with the enhanced inhibitory effect on growth, the co-exposure to isosteviol and duloxetine showed the largest variation in the end point mutant library composition (Fig. 4b).

To define gene hits, we set an absolute log2 fold change cutoff of 0.5 for conditional fitness, which is the proportion of gene mutants at the endpoint of chemical perturbation compared to the DMSO control. Thirteen mutants responded to co-exposure, and five to duloxetine (Fig. 4c, supplementary table 15). The strongest conditional negative fitness (loss of mutants) maps to genes encoding membrane proteins, suggesting their key role in survival during co-exposure. These include a putative magnesium transporter of the CorA family (NQ542_03545), a putative hydrophobic compound transporter of the Fadl family (NQ542_08455), a non-essential domain of the outer membrane protein assembly factor BamD (NQ542_03840) and a putative c-di-AMP phosphodiesterase (NQ542_10115) (supplementary figure 11). NQ542_10115 is an ortholog of the Pde c-di-AMP regulator implicated in cell membrane homeostasis in *Porphyromonas gingivalis* (50% identity, 94% coverage)^47^. When comparing Tn-Bar-Seq gene fitness score with the proteomics data, *corA*-family transporter NQ542_03545 stands out as a candidate with strong negative fitness and increased protein abundance both for duloxetine and co-exposure (supplementary figure 11). However, magnesium supplementation did not subvert compound exposure phenotype in *P. merdae* (supplementary figure 12) suggesting this transporter has alternate, yet unknown, role in cellular homeostasis.

The mutant for putative transcriptional regulator *acrR* (NQ542_01170), which is an ortholog of an efflux regulator in *Phocaeicola vulgatus*^*48*^, exhibited negative fitness in accord to chemical transport changes (supplementary figure 11). While above-mentioned genes were also significant hits for duloxetine alone, their negative fitness was further exacerbated when isosteviol was present. Among the other top gene hits are genes from central metabolism and vitamin synthesis, which are often observed for xenobiotics having a mode of action different to canonical antibiotics^49^. Overall, both approaches confirm that membrane integrity and transport are crucial for survival under the duloxetine-isosteviol combination.

### Isosteviol and duloxetine alter bioaccumulation

Previously we observed that duloxetine bioaccumulation can alter metabolism in gut bacteria^33^. We therefore assessed whether the synergistic effects of duloxetine and isosteviol are reflected in, or contributed to, by the differences in isosteviol and duloxetine bioaccumulation by *P. merdae* and *R. intestinalis*. For comparison, we also included *Segatella copri*, a bacterium that shows significant inhibition by duloxetine in monoculture and a marked decrease in the community when treated with either duloxetine alone or co-treated with isosteviol. While *S. copri* bioaccumulated both compounds to the similar extent under all tested conditions, *R. intestinalis* bioaccumulated isosteviol only when duloxetine was present (supplementary figure 13, supplementary tables 16 and 17). On the other hand, *P. merdae* bioaccumulated isosteviol in the absence of duloxetine. The altered bioaccumulation is consistent with the altered metabolite transport suggested by the proteomic and genetic analyses and likely contributes to the synergy between isosteviol and duloxetine.

### Duloxetine and isosteviol exposure affect secreted metabolites

Genetic analysis and altered bioaccumulation both indicated changes to cross-membrane transport of small molecules. We therefore hypothesized that co-exposure of duloxetine and isosteviol alter the metabolite secretome. To test this, we used untargeted liquid chromatography mass spectrometry (LC-MS) to characterize secreted metabolites as well as whole culture metabolic profile (cells and supernatant) of *S. copri, R. intestinalis* and *P. merdae* (Methods, supplementary tables 18-23).

The three tested species showed different response to the compound exposures (Fig. 5a-c, supplementary figure 14). For *S. copri*, both the whole culture and supernatant metabolites clustered by treatment, indicating a co-treatment effect on the metabolome (Fig. 5a). In monoculture, no combinatory effect was observed; however, community growth was reduced by both single compounds and co-exposure compared to DMSO. Thus, the altered metabolism may contribute to the decreased competitiveness of *S. copri* in the microbial community (Fig. 2e, supplementary figure 6).

The metabolites of *R. intestinalis* (both, whole culture, and supernatant) clustered by duloxetine treatment (Fig. 5b). The impact of isosteviol was minor and similar to that of the DMSO control, reflecting the changes observed in protein abundances wherein the effect of duloxetine was dominant compared to that of isosteviol (Fig. 3c). The combinatory growth effects in *R. intestinalis* may thus be mainly due to duloxetine-dependant alterations in isosteviol bioaccumulation (supplementary figure 13).

In the case of *P. merdae*, whole culture metabolites clustered based on the corresponding treatments. Yet, for the supernatant, all treatments cluster together, separate from the vehicle (DMSO) (Fig. 5c, supplementary figure 14c), albeit with a slight separation of co-treated samples from DMSO. The similar secretion profiles for both mono- and co-exposure suggests the key role of altered abundance of membrane proteins and transporters as revealed by proteomics analysis (Fig. 3c).

To gain indications on the nature of the impacted metabolites, we selected 19 peaks from the untargeted analysis that showed significantly different abundance in the three microbes treated with duloxetine and isosteviol as compared to the vehicle treatment. Of the 19 peaks, sixteen could be annotated as di- and tripeptides based on matching of MS2 spectra to an online database using the PEP search tool^50^ (supplementary figure 15, supplementary table 24, Methods). For all three species, both increase and decrease of di- and tripeptides as a response to the combined treatment were observed. Microbial di- and tripeptides have been shown to have antihypertensive properties^51^ and to be involved in *Listeria monocytogenes* infection^52^ and intestinal inflammation^53,54^. In conclusion, the identified di- and tripeptides, which showed altered abundance in response to duloxetine and isosteviol treatment, may influence various biological processes and modulate the host’s response to microbial presence.

### Co-exposure impacts primary metabolism in *R. intestinalis* and *P. merdae*

To further pinpoint the metabolic effects of isosteviol and duloxetine, we used targeted LC-MS analysis to quantify 46 metabolites (32 amino acids and derivates, 10 organic acids and 4 short chain fatty acids) in *S. copri, R. intestinalis* and *P. merdae* (supplementary tables 25-27). For *S. copri*, three metabolites showed significant (p<0.05) changes in response to the duloxetine-isosteviol combination; cadaverine, nicotinic acid and pyridoxal (supplementary figure 16, 17, 18). *R. intestinalis* showed changes in glutamine, butyric acid and isovaleric acid. Notably, glutamine production increased by circa 50% and butyric acid decreased by more than 25% upon co-exposure to duloxetine and isosteviol (Fig. 5d, supplementary figure 17, 18, 19).

In absolute terms, the concentrations of glutamine and butyric acid changed substantially, by circa 65 µM and 0.6 mM, respectively, thus implying relevance for microbe-host interaction. Butyric acid has previously been associated with anti-inflammatory properties, increased insulin sensitivity, and protection against diet-induced obesity^55–57^, and these effects have also been linked to *R. intestinalis*^*58–60*^. In contrast, increased isovaleric acid concentrations in faeces have been shown to correlate with depression^61^. In *P. merdae*, compound co-exposure led to significant changes in agmatine, asparagine and pyridoxamine concentrations. While total concentrations of agmatine and pyridoxamine are in the lower µM range, the concentration of asparagine changed by >200 µM indicating substantial changes in amino acid metabolism (Fig. 5e, supplementary figure 20, 17, 18). The altered peptide concentration uncovered by the untargeted analysis further highlights the impact on amino acid metabolism. Together, synergistic effects of isosteviol and duloxetine on gut bacteria alter secretion of metabolites linked to host health.

### Altered microbial metabolism modulates toxicity and cytokine secretion by host cells

Gut bacterial metabolites are key mediators of microbiota-host interaction^62^. We therefore next examined whether metabolic changes caused by the isosteviol-sweetener exposure impact host cells. We first tested the toxicity of supernatants from the 25-member gut bacterial community (Fig. 2) exposed to isosteviol, duloxetine, and their combination on HeLa cells. HeLa cells are commonly used in cell biology and here serve as a starting point to assess potential effects of an altered secretome.

Community supernatant was added to the cell growth medium (DMEM) for 24 hours before cell viability was assessed (Methods). We observed a significant increase in toxicity to HeLa cells in the supernatant of communities treated with the combination of isosteviol and duloxetine, compared to the solvent-only control, DMSO (Fig. 6a, supplementary table 28 and 29). To exclude the direct effects of the compounds on HeLa cells, we included an additional control: the DMSO control plus 50 µM isosteviol and duloxetine added after harvesting the supernatant. Both, the DMSO and the DMSO plus compounds control (drug control) were significantly less toxic than the supernatant of the community treated with the isosteviol-duloxetine combination.

To assess the impact of altered secretome in more physiologically relevant cells, we used colon carcinoma-derived Caco-2 cell lines. In addition to cytotoxicity, we also assessed the secretion of 13 cytokines. Cytokines are soluble proteins secreted by immune cells and intestinal epithelial cells (IEC) to mediate intercellular communication in homeostasis but also play a relevant role during inflammation^63,64^. IEC-derived cytokines such as interleukin-6 (IL-6), interleukin-8 (IL-8) and C-X-C motif chemokine ligand 10 (CXCL10 also known as IP-10), are key indicators of inflammation-associated damage in the gut in response of bacterial infection and Inflammatory Bowel Disease^65–68^. A reduction in these cytokines might indicate an anti-inflammatory response; however, it could also indicate the gut’s impaired ability to combat pathogens effectively^69–71^.

While no excess cytotoxicity was observed compared to the drug control (Fig. 6b, supplementary figure 21), the secretion of three cytokines was substantially modulated: IL-6, IL-8 and CXCL10. All three cytokines were stimulated by the supplementation of 20 % community supernatant to the culture medium. The supernatant of a community co-treated with isosteviol, and duloxetine (50 µM each) promoted the secretion of IL-6 and IL-8 to a much lesser extent but did not affect that of IP-10 (Fig. 6c-e). IL-6 and IL-8 are key regulators of neutrophil biology: IL-6 promotes neutrophil production in the bone marrow, while IL-8 acts as a potent chemoattractant for neutrophils. Both cytokines exhibit both, pro- and anti-inflammatory effects, with their roles depending on the specific biological context^72,73^. They were previously shown to be stimulated by the presence of butyrate, LPS, IL-1beta and TNFalpha^68,74,75^. The community supernatant of the 25 bacteria community contains ∼3 mM butyric acid and 2 mM propionic acid. Both SCFAs are significantly reduced in the community supernatant following treatment with duloxetine and isosteviol (supplementary figure 22). Although Caco-2 did not produce IL-1beta and TNFalpha (supplementary table 30), changes in SCFA production, in combination with microbial molecules such as LPS and flagellin, may contribute to IL-6 and IL-8 secretion by these cells. The addition of LPS and a combination of propionate and butyrate, respectively, was sufficient to induce secretion of IL-8 and IP-10 (supplementary figure 23).

## Discussion

Previous cohort and animal studies have linked sweetener consumption to the microbiome-level changes, yet connecting most of these effects mechanistically to microbial action has not been realised to date. Our data opens a mechanistic window into the sweetener-xenobiotic-microbe-host interactions by bringing forward their impact on bacterial growth and secreted metabolites. As the latter is one of the main pillars of microbiome-host interactions^62,76^, it is likely that some of the reported effects of sweeteners and sweetener-xenobiotic combinations on the host are through changes in bacterial metabolites. Our study identified metabolites with known function in the microbiome-host interaction such as butyrate and found that sweeteners can alter the impact of bacterial metabolites on host cells. The study also opens paths for future mechanistic work. For example, as there are inter-individual differences in microbiota at strain level^77^, it will be important to investigate whether the metabolic effects observed in our study are strain specific. Further, extensive survey of *in vivo* concentrations of relevant compounds and their kinetics and dynamics within the host will be needed for better connecting *in vitro* data with cohort studies.

Emergent effects due to mixture of compounds, the so-called ‘mixture effects’ or ‘mixture model’, is commonly considered in the context of drug efficacy and toxicity^78^. In the context of gut microbiota, antagonistic interactions have been linked to antimicrobial resistance^26^. However, emergent effects of chemical mixtures beyond antibiotic resistance remain sparsely studied and underappreciated. Our results emphasize the need to study mixture effects more broadly, especially for food ingredients like sweeteners that are consumed daily by millions. Our multi-omics approach spanning genetic, proteomic, and metabolomic analyses underscore the need to study physiological effects beyond growth and provide a starting point to systematically unravel the complex sweetener-microbiome-host interactions.

## Methods

### Microbe selection and growth conditions

The 25 human gut microbial bacteria, 24 species and two different strains of *E. coli*, were selected based on abundance in the human gut and to cover a broad phylogenetic and metabolic diversity representative of the healthy microbiota, including also potential pathogens and probiotics (supplementary table 2)^28,33^. Unless otherwise indicated, all bacteria were grown as liquid cultures in Gifu Anaerobic Medium broth, modified (mGAM, HyServe, Germany), an undefined rich medium that has been successfully used in cultivating gut microbial communities and individual species in previous screening studies^33,79^. And has further been demonstrated to support the growth of most gut bacteria strains in monoculture^28^. Culturing was carried out in a vinyl anaerobic chamber (COY) at 37 °C with 12 % carbon dioxide, 86 % nitrogen and 2 % hydrogen. All experimental cultures were started from the second passage culture after inoculation from a glycerol stock. All media, buffer, glass and plasticware used in the study were exposed to the anaerobic conditions at least 12 hours before use.

### Plate preparation and screening of sweeteners and combinatory compounds

Screening was done as previously described in Müller et al and Maier et al^79,80^. All tested sweeteners and compounds were dissolved in dimethyl sulfoxide (DMSO). Final screening concentration of all compounds was 50 µM (an amount estimated to reach the gut on average, supplementary table 3b) in mGAM. The total screening volume used was 100µL per well. The final concentration of the DMSO solvent was kept constant at 1 % in both, single compound and combination screens. The starting OD_600_ was adjusted to 0.05 for all strains tested. Plates were sealed with breathable membranes (BreatheEasy®) to allow gas exchange. Optical density readings were taken at 600 nm, using the BioTek Epoch 2 Microplate Spectrophotometer and BioStack 4 Microplate Stacker (both from Agilent) fitted in a custom-made incubator system (EMBL mechanical workshop) inside the anaerobic chamber. The screen was carried out under anaerobic conditions in 96-well plates and OD_600_ was measured every hour for 24 hours. Each plate had six negative controls (bacteria with an equivalent amount of DMSO). Positive and negative effects of sweeteners on bacterial growth were assessed using respective negative controls on each plate. Each strain was screened at least three times (biological replicates) with two replicates per plate (technical replicates), adding up to 6 replicates in total.

**Retesting and dose response** was done as described above, in mGAM and 1 % DMSO and growth was monitored hourly for 24 hours. All 33 bacteria - sweetener interactions were retested in the sweetener concentrations 25, 50, 100, 200 and 400 µM. We regarded the retests as verified if two or more concentrations revealed to be significant (Welch’s t-test, N>3, p<0.05, supplementary tables 31a and b).

### Data processing and growth quantification

Optical density readings were analysed by plate for each strain and replicate. Growth curves were fitted to each well using the ‘Growthcurver’ package in R^81^. The growth of each microbe in presence or absence of compounds was quantified using area under the curve (AUC) of the fitted growth equation. The entire screen was quality checked by overlaying all fitted and raw curves of technical and batch replicates for each species. Out of 39,313 screened samples, 492 noisy wells and 136 failed fits (<2% in total) were removed from further analysis, making a total of 38,685 samples entering further analysis (see supplementary table 4a for the complete dataset including the excluded wells).

### Data normalisation and hit selection

Non-systematic variations were controlled by normalising data within each plate to have comparable results across replicates. The wells were normalised by using 6 control wells present on each plate. Median percent inhibition (MPI) was used to normalise samples: MPI = sample AUC/ median (controls AUC). Welch’s t-test was performed between all replicates (two technical and 3 biological replicates) of each sample per compound and their corresponding control wells. A two-sided p-value at an α-level of 0.95, standardised effect sizes and log2 of average fold changes were calculated. The p-values were adjusted for multiple hypothesis correction using the ‘Benjamini-Hochberg’ method. Also, log_2_ of average fold change (= normalised AUCs) between technical replicates was reported for each batch replicate. Finally, the hit compounds were selected using two criteria 1) p-value < 0.05 and 2) log_2_FC of at least 20% (see supplementary tables 4 and 5 for p-values and single and combinatory screen details).

### Determination of combinatory interactions

To determine interactions between sweeteners with the combinatorial compounds the Bliss model of independence was used^40^, which divides interactions in synergistic, antagonistic or no interaction. Here we used the log-transformed normalised AUC as surviving fraction of microbes as developed by Demidenko et al^41^. According to this model, two compounds (A and B) act independently, if the surviving fraction of cells upon simultaneous administration of both the compounds equals the product of compounds given separately. A synergy or an antagonism exist if SF_AB_ < S_A_S_B_ or SF_AB_ > S_A_S_B_ (supplementary figure 3a). A two-sided p-value was computed for determining synergy or antagonism and an interaction was seen as significant at a p-value threshold of < 0.05 and a synergy or antagonism of at-least 20% as compared to the control.

### Synthetic microbial communities

Synthetic microbial communities were assembled from the 25 gut bacteria strains and species that were used in the growth screen in monoculture. Each bacterium was grown for two passages as described above and combined with all others in equal proportions, based on optical density under anaerobic conditions. The species mix was then diluted 1:20 with fresh mGAM medium alone or mGAM containing 50 µM of the compounds dissolved in DMSO (0.2% total DMSO, also in the negative control). Passaging was done every 24 hours.

### 16S rDNA sequencing and data analysis

To ensure that separation on species level is feasible, microbial species had been selected based on distinguishable V4 rDNA sequence prior to *in vitro* community assembly. All samples for community compositional analysis were examined by targeting the fourth hypervariable (V4) regions of the 16S rDNA gene as predicted by barrnap (Version 0.9). DNA was extracted using the MagMAX™ Microbiome Ultra Nucleic Acid Isolation Kit along with the KingFisher Flex Purification System. Library preparation for the 16S V4 was done by the European Molecular Biology Laboratory genomics core facility using a customized barcoding, amplification, and purification approach. The barcoded amplicons were pooled in equal concentrations and sequenced on the Illumina MiSeq platform using 2 × 250 base pairs at the EMBL genomic core facility in Heidelberg, Germany. Analysis of amplicon sequencing data was done as previously described in Blasche et al^82^.

### Preparation of *P. merdae* and *R. intestinalis* samples for protein expression analysis

Both species were inoculated from frozen stock cultures, three replicates each, into mGAM medium and passaged twice over night. Passage two was transferred into 60 ml (120 ml for *R. intestinalis*) medium and pre-exposure0 samples were collected directly hereafter (three per species, one sample per replicate, three in total). The second pre-exposure samples were collected upon reaching exponential growth (after three hours for *R. intestinalis* and 2.5 hours for *P. merdae*), directly prior to splitting each replicate into four equal sized batches, one per condition: Mock (DMSO, 0.2 %), 50 µM duloxetine, 50 µM isosteviol and a combination of both, 50 µM each. This resulted in 12 treatment samples, three per treatment, whereas each of the replicates could be traced back to one preculture. Bacteria were grown in the presence of the compounds at 37°C for four hours under anaerobic conditions, before final sample collection. Pre-exposure samples were collected as controls to monitor changes in protein expression in different growth stages and thus to minimise the capturing of expressional differences due to altered in microbial growth (tested compounds affect species growth). For collection, samples were centrifuged at 4000 rpm for 25 min at 4°C, 15 ml PBS were added to the pellet followed by centrifugation at 4000 rpm for 10 min. Pellets were heat-inactivated in mass spectrometry-compatible lysis buffer containing 100 mM triethyl ammonium bicarbonate (TEAB) and 0.05% RapiGest (Waters, Wilmslow), twice by heating at 80°C for 10 min followed by cooling for 10 min on ice. The inactivated cells were stored at -80°C until further processing.

### Proteomics sample processing and data analysis

The heat inactivated cells were lysed by sonication to release the proteins and equal concentration of proteins from each sample were taken for tryptic digestion. Towards this, the proteins were denatured by treatment with 0.1% RapiGest at 80 °C for 10 min, reduced with 4 mM dithiothretol (DTT), alkylated with 14 mM iodoacetamide (IAA), and subsequently digested overnight at 37 °C with trypsin at 1:50 protease:protein ratio. Equal concentration of the digested peptides was taken for TMTpro 18plex labelling as per the manufacturer’s instructions. In brief, 35 ug of tryptic peptides from samples were labelled with respective TMT reagents at 1:10 peptide: label ratio in 50 uL of 100 mM TEAB pH 8.0 buffer. The TMTpro labels were randomized across the samples to be compared. The labelling mix was incubated at 21 °C for 1 h with intermittent vortexing and the reactions were quenched with hydroxylamine at 0.4% final concentration. Equal volumes of the labelled peptides were pooled to multiplex and vacuum dried. The dried peptides were resuspended in 100 µM TEAB containing 1% trifluoro acetic acid (TFA) and desalted using C18 spin columns (Pierce) as recommended by the manufacturer. Prior to desalting RapiGest was hydrolysed and removed along with any insolubles. The multiplexed samples were subsequently fractionated offline by HpH reverse phase chromatography into 12 concatenated fractions. The fractions were dried and resuspended in 0.1% TFA in 3% Acetonitrile (ACN) for mass spectrometry.

An equal volume of peptides from each fraction were analysed on an Orbitrap Eclipse™ mass spectrometer coupled in-line to an Ultimate 3000 RSLC™ nano LC system. The peptides were loaded onto a trap column (PepMap100, C18, 300 μm X 5 mm) and separated on an analytical column (Easy-Spray C18 75 µm x 500 mm 2 µm) at 300 nL/min flow rate using 0.1% Formic acid (FA) in water and 0.1% FA in 80% ACN as mobile phases A and B, respectively. The gradient program involved raising %B 3-25 over 140 min, followed by further increasing to 40% B in 16 min. The overall inject-to-inject run time was 184 min.

Reporter ion quantification was carried out from MS3 fragmentation spectra acquired employing a synchronous precursor selection (SPS) based on real time search (RTS) workflow. Uniprot protein fasta databases UP000004276 (unreviewed TrEMBL db downloaded on 20230718, contains 4351 seq) and UP000004828 (unreviewed TrEMBL db downloaded on 20220411; contains 4697 seq) were configured as databases for RTS for *P. merdae* and *R. intestinalis* datasets, respectively. The MS1 precursors were scanned in 415-1500 mass range at 120000 resolution in the orbitrap mass analyzer. Gas phase fractionation was employed by setting 3 concurrent compensation voltages (−45, -60, -75) in FAIMSPro. At each compensation voltage, the precursors with a charge state of 2-5 and a minimum intensity of 5e3 were isolated in quadrupole at 0.7 isolation width, fragmented by HCD at normalized collision energy (NCE) 32, and analysed in ion trap mass analyzer under turbo mode. Top 10 fragments retrieving a match in 35ms on real time search were synchronously selected for MS3 fragmentation by HCD at 55 NCE, and the reporter ion spectra was scanned in 100-500 mass range at 120000 resolution in orbitrap. Max injection time was set at 246ms. Cys carbamidomethylation and TMTPro on Lys and N-terminus were given as fixed modifications and Met oxidation as variable modification for real time search. Decoy search and FDR filtering were enabled.

The raw data were imported as fractions into Proteome Discoverer 3.0 and analysed against respective protein databases using a predefined workflow employing both SequestHT and Comet search algorithm. Precursors and fragments matched with max 10 ppm and 0.6 Da mass tolerance, respectively. Two missed cleavages were tolerated and TMT labelling on Lys and N-terminus and Carbamidomethylation of Cys were set as fixed modification and oxidation of Met as variable modification. Common contaminants matching to cRAP database from GPM were filtered out from search results before empirical analysis. Matches to a decoy database was utilised for False Discovery Rate (FDR) validation employing Percolator and matches passing 1% FDR threshold were taken further for defining qualitative and quantitative identification. Quantification was carried out based on S/N values of reporter ions of the identified peptides that are unique to the representative proteins. The protein abundance was enumerated by summing the abundances of peptide groups matching to respective proteins. Normalization factor was generated with respect to the summed abundances of all the identified proteins and the TMT channel with the highest total protein abundance was used as the reference channel. The normalized abundance values were used for quantitative comparison across sample sets.

### Tn-Bar-seq with *P. merdae* genome-wide transposon library

Two vials of the pooled library of *Parabacteroides merdae* ATCC 43184 mapped to GenBank: CP102286.1 (Voogdt et al., unpublished) were thawed and used to inoculate mGAM with 30 μg/mL erythromycin and grown until mid-exponential phase. At this stage, the culture was used to inoculate at OD_600_ 0.02 1 mL assay cultures in a deep well plate, with mGAM containing 50 μM of duloxetine, isosteviol alone or in combination in duplicates and triplicate for the DMSO control. The libraries were grown until early stationary phase at 37° C in an anaerobic chamber, where end point samples were taken for gDNA extraction. The growth of the mutant library was monitored by growing in parallel a 100 μl aliquot of the experiment and measuring OD_600_ in the same set up than the initial growth screen (BioTek Epoch 2 Microplate Spectrophotometer and BioStack 4 Microplate Stacker). gDNA was extracted with MagMAX Microbiome Ultra Nucleic Acid Isolation Kit and the barcodes amplified with NEB Q5 Hot Start HF X2 with custom Tn-Bar-Seq primers. The amplicon with the barcodes were sequenced with Illumina NextSeq 75 nt SE. Barcodes were counted with 2FAST2Q^83^ and changes in mutant gene abundance were analysed with TRANSIT using reads in the central 80% of the open reading frame and normalised by Trimmed Total Read-count and compared to the DMSO control by resampling set at 20000^84^. To cross-compare Tn-Bar-Seq to proteomics, annotations were matched by BlastP best reciprocal hit, Protein Accession to Locus ID match can be found at (supplementary table 8).

### Bioaccumulation of duloxetine and isosteviol

Bacteria were inoculated from second passage at a starting OD_600_ of 0.05 and grown anaerobically at 37°C in the presence of the compounds and the combination. Controls, the solvent control (bacteria plus DMSO) and the incubation control (bacteria-free mGAM with the compounds), were treated the same way. The incubation control tests for bacteria-independent compound degradation and, when compared to the whole culture, may reveal compound metabolism if a significant amount is metabolized within the observed time frame. After 24 hours the sampled were taken out and whole culture (wc, no further treatment), supernatant (sup, centrifuged at 4000g for 25 min) and compound control were collected and stored at -80 ºC until extraction.

### Sample extraction for untargeted metabolite analysis and compound bioaccumulation

LC-MS samples, both whole cultures and supernatants, were extracted by adding 4x the amount of ice cold (−20°C) extraction buffer (1:1 ratio of methanol and acetonitrile with internal standards, 75µM caffeine and 60µM ibuprofen, spiked in) followed by an incubation at 4 ºC for 15 min. Samples were then centrifuged at 4 ºC and 4000 rpm for 25 min and supernatants were transferred to 96-well plates for LC-MS analysis. Samples for concentration calibration (1 in 2 dilution of compounds) and bacteria-free compound controls were processed in the same way.

### Reverse phase LC-MS for untargeted metabolite analysis and compound bioaccumulation

LC-MS analysis was performed on an Agilent 1290 Infinity II LC system coupled with an Agilent 6546 Q-TOF with JetStream ESI source. The separation was performed using a ZORBAX RRHD Eclipse Plus column (C18, 2.1 × 100 mm, 1.8 μm; Agilent 858700-902) with a ZOBRAX Eclipse Plus guard column (C18, 2.1 × 5 mm, 1.8 μm; Agilent 821725-901) at 40ºC. The multisampler was kept at a temperature of 4 ºC. The injection volume was 1 μL and the flow rate was 0.4 mL/min. The mobile phases consisted of A: water + 0.1% formic acid + 5 mM ammonium formate; B: methanol + 0.1% formic acid + 5 mM ammonium formate. The 10 min gradient started with 5% solvent B, which was increased to 90 % by 5 min and then further increased to 100 % by 7 min, before returning to 5 % solvent B for a 3 min re-equilibration. The instrument was operated in TOF MS acquisition mode in either positive or negative polarity. (30-1500 m/z). The source parameters were as follows: gas temperature: 200ºC, drying gas: 9 L/min, nebulizer: 20 psi, sheath gas temperature: 400 ºC, sheath gas flow: 12 L/min, VCap: 3000 V, nozzle voltage: 0 V, fragmentor: 110 V, skimmer 45 V, Oct RF Vpp: 750 V. The online mass calibration was performed using a reference solution (positive: 121.05 and 922.01 m/z; negative: 112.99 and 1033.99 m/z). Collision energies used were 0 V, 10 V, 20 V, 40 V. Compounds of interest were identified based on their retention time, accurate mass, and fragmentation patterns, using pure standards for method development, compound identification and calibration.

**Untargeted metabolite analysis** was performed using the Agilent MassHunter Workstation Profinder 10.0 software. Area under the curve was extracted, using the Batch Recursive Feature Extraction function for small molecules and peptides. Retention time was restricted from 0.5 to 8.0 minutes and a tolerance was set to 0.05 minutes. Peaks above a raw hight of 4000 in at least one sample were extracted and peaks below 4000 in all samples were regarded as noise and discarded.

### Untargeted metabolomics peak identification

Raw data files were converted to .mzML format using msconvert (ProteoWizard) with peakPicking and zeroSamples functions applied. The .mzML files were uploaded to R and XCMS used to perform peak picking, refinement, correspondence, and peak filling with the parameters listed in supplementary table 32^85^. Statistical analysis was performed using MetaboAnalyst 6.0^86^. Near-constant variables were filtered by interquartile range so 25 % of features were removed and the data were then normalized by sum, log transformed and pareto scaled. Correlation analysis was performed using the PatternHunter tool to highlight features more significantly altered by combination treatment with both duloxetine and isosteviol than by either treatment alone. The correspondence of feature intensity to this pattern was determined using Pearson correlation. A significance threshold of correlation coefficient > ± 0.8 and p-value < 0.05 was employed. Metabolite identification was performed by comparison of accurate mass to theoretical mass with an error tolerance of ±10 ppm and where possible by matching of MS2 spectra to database MS2 spectra using the PEP Search tool^50^.

**Targeted metabolomics** samples were prepared by adding 120 μl of extraction buffer (1:1 methanol:acetonitrile, 0.1% formic acid, 20 μM caffeine, amoxicillin and donepezil as internal standards) to 80 μl of sample, followed by a 30 min incubation at 4ºC and centrifugation (5min, 4ºC, 3,200g) to remove precipitated proteins and salts. For amino acid and organic acids (except SCFA) analysis, samples were diluted 1:10 in water prior to extraction. Analysis was performed on an Agilent 1290 Infinity II LC system coupled with an Agilent 6470 triple quadrupole mass spectrometer with JetStream ESI source operated in dynamic multiple reaction monitoring (dMRM) mode. Amino acids were measured using a low pH HILIC as described previously^87^. In brief, 0.25 μl of sample was injected and analytes were separated on a Waters Acquity BEH Amide column (1.7 µm, 2.1 mm x 100 mm) maintained at 35ºC. The gradient used a constant flow rate of 0.9 ml/min and proceeded as follows: 0 min: 15% buffer A (1:1 acetonitrile:water, 10 mM ammonium formate, 0.176% formic acid) and 85% buffer B (95:5:5 acetonitrile:methanol:water, 10 mM ammonium formate, 0.176% formic acid), 0.7 min: 85% B, 2.55 min: 5% B, 2.9 min: 5% B, 2.91 min: 85% B, 3.5 min: stop time.

Organic acids were measured using a high pH HILIC method as described previously^88^. In brief, 2 μl of sample was injected and analytes were separated on a Waters Atlantis Premier BEH Z-HILIC (1.7 µm, 2.1 mm x 100 mm) column maintained at 35ºC. The gradient used a constant flow rate of 0.5 ml/min and proceeded as follows: 0 min: 15% buffer A (15mM ammonium bicarbonate in water, pH 9) and 85% buffer B (15mM ammonium bicarbonate in 9:1 acetonitrile:water, pH 9), 1 min: 85% B, 3 min: 25% B, 4 min: 25% B, 4.01 min: 85% B, 5 min: stop time.

SCFA were measured using a porous graphitic carbon column as described previously^89^. In brief, 1 μl of sample was injected and analytes were separated on a Thermo Scientific Hypercarb PGC column (3 µm, 50 mm × 2.1 mm) maintained at 40ºC. The gradient used a constant flow rate of 0.15 ml/min and proceeded as follows: 0 min: 100% buffer A (water, 0.1% formic acid) and 0% buffer B (acetonitrile, 0.1% formic acid), 4 min: 60% B, 4.1 min: 100% B, 6 min: 100% B, 6.01 min: 0% B, 9 min: stop time.

A pooled QC sample, blanks and serial dilutions of a mixed analytical standard were injected throughout each run. Data were analysed using MassHunter Workstation Quantitative Analysis for QQQ v10.1. Concentrations were obtained via external calibration using standard curves.

### Cytotoxicity assay

HeLa and Caco-2 cell lines (passage numbers 10 and 15, respectively) were obtained from the Cambridge University Biological Services Unit, authenticated by ATCC and tested for Mycoplasma before use. The cells were cultured in DMEM (Gibco), containing 10% Fetal Bovine Serum (FBS; Sigma-Aldrich) in standard conditions (37 °C, 5% CO2). Cells were trypsinized and seeded in 96-well plates at a density of 2.5 × 10^4^/well, in 80 μL of phenol-red and FBS-free media, allowed to adhere overnight and treated by adding 20 μL of community spent media for 24 h. Untreated cells were used as negative control taken as 100% viability, further negative controls include supernatant of DMSO treated microbial community with (DMSO control) and with drugs added prior to cell culture exposure (drug control), see Fig. 6a and b. All microbial supernatants were centrifuged and sterile filtered (Millex PVDF syringe filter, pore size 0.22 μm, Merck) prior to application. 5 μM staurosporine was used as positive control for cytotoxicity induction. Cytotoxicity was assessed by adding 20 μL of CellTiter-Blue Reagent (Resazurin) (Promega) and incubated at 37 °C, 5% CO^2^ for 3 hours. Fluorescence was measured using a VarioSkan (Thermo) plate reader, at 560/590 nm.

### Cytokine assay

Caco-2 cells were seeded and treated as described for the cytotoxicity assay. After 24 h of treatment, supernatants were harvested and stored at -80 °C until use. For cytokine assessment, LEGENDplex™ Human Essential Immune Response Panel (13-plex) (BioLegend Cat. No. 740930) was used. In brief, 25 μL of the supernatant was mixed with capture beads coated with antibodies specific to every target cytokine. After 2 h incubation at RT, samples were washed and 25 μL of detection biotinylated antibodies were added and incubated for 1 h RT. Finally, 25 μL of streptavidin-PE conjugated was added to every sample and washed after 30 min of RT incubation. A BD LSRFortessa™ was used for data acquisition using the settings specified by the supplier. Data was analysed using BioLegend’s LEGENDplex™ data analysis software.

### Supernatant titration for HeLa cytotoxicity assay

To determine what proportion of cell culture medium (DMEM) can be replaced by bacterial community supernatant, we tested three percentiles of supernatant in HeLa cells: 5, 10 and 20%. Although the supernatant decreased cell viability with increasing concentration, even the highest concentration of 20% spent medium revealed a cell viability of 84%, high enough to be suitable for testing (supplementary figure 24)

### Staurosporine and LPS titration

The amount of staurosporine to be added for apoptosis induction was determined for HeLa and Caco-2 by titration (supplementary figures 25 and 26). Induction of cytokines by staurosporine and LPS was titrated in Caco-2 cells. The amount of staurosporine used to induce apoptosis (>4 µM) is higher than the one inducing cytokine secretion (supplementary figure 23).

## Replicates and statistical tests

Technical replicates refer to replicates within the same batch (for example, two bacterial cultures inoculated from the same starting culture or on the same plate), whereas biological replicates refer to independent experiments. All indicated sample numbers (or displayed data points) refer to distinct samples (biological and technical replicates) and not to repeated measurements of the same samples. All statistical tests are two-sided. FDR corrections were performed using the Benjamini–Hochberg procedure. If not stated otherwise, statistical significance was determined using student’s t-test.

## Code and Data availability

Proteomics data is available at Pride database (https://www.ebi.ac.uk/pride/) under ID PXD045140 (reviewer access available with the username: and password: SYdQthxG, please login on left side!). Codes used for processing and raw screening data are available at https://github.com/periwal45/SW_SF100. Tn-Bar-Sequencing data and 16S amplicon sequencing data are deposited at ENA database (https://www.ebi.ac.uk/ena/browser/home) with accession ID PRJEB76554 and PRJEB76553, respectively. Metabolomics data is deposited at MetaboLights database under the ID MTBLS11480.

## Author contribution

SB, VP and KRP conceived the study and planned the overall experiments. VP and SB designed and performed the screening experiments and community assays. VP analysed the data from the screening assays. SB, RG and VP analysed the data from the community assays. BR and SB designed, performed and analysed data from the proteomics experiments. IR performed and analysed *P. merdae* genome wide mutant library screens. SB and AL designed, performed, and analysed bioaccumulation and metabolomics experiments. RB identified metabolites in untargeted metabolomics data. NBC and SB designed and performed the toxicity assays in HeLa and Caco-2 cells. SK, RG and SB designed, performed and analysed data from the targeted LC-MS experiments. SB, VP and KRP wrote the manuscript. All authors read and/or commented on the manuscript.

## Acknowledgements

We thank the EMBL Gene Core facility for help with the 16S amplicon sequencing. This project has received funding from the European Union’s Horizon 2020 research and innovation programme (grant nos. 866028 and 814408) and from the UK Medical Research Council (project no. MC_UU_00025/11).

## Competing interests

Authors declare no competing interests.

## Notes

### Competing Interest Statement

The authors have declared no competing interest.

