## Supplementary figures for "Impact of low-calorie sweeteners on gut bacteria is modulated by common xenobiotics"

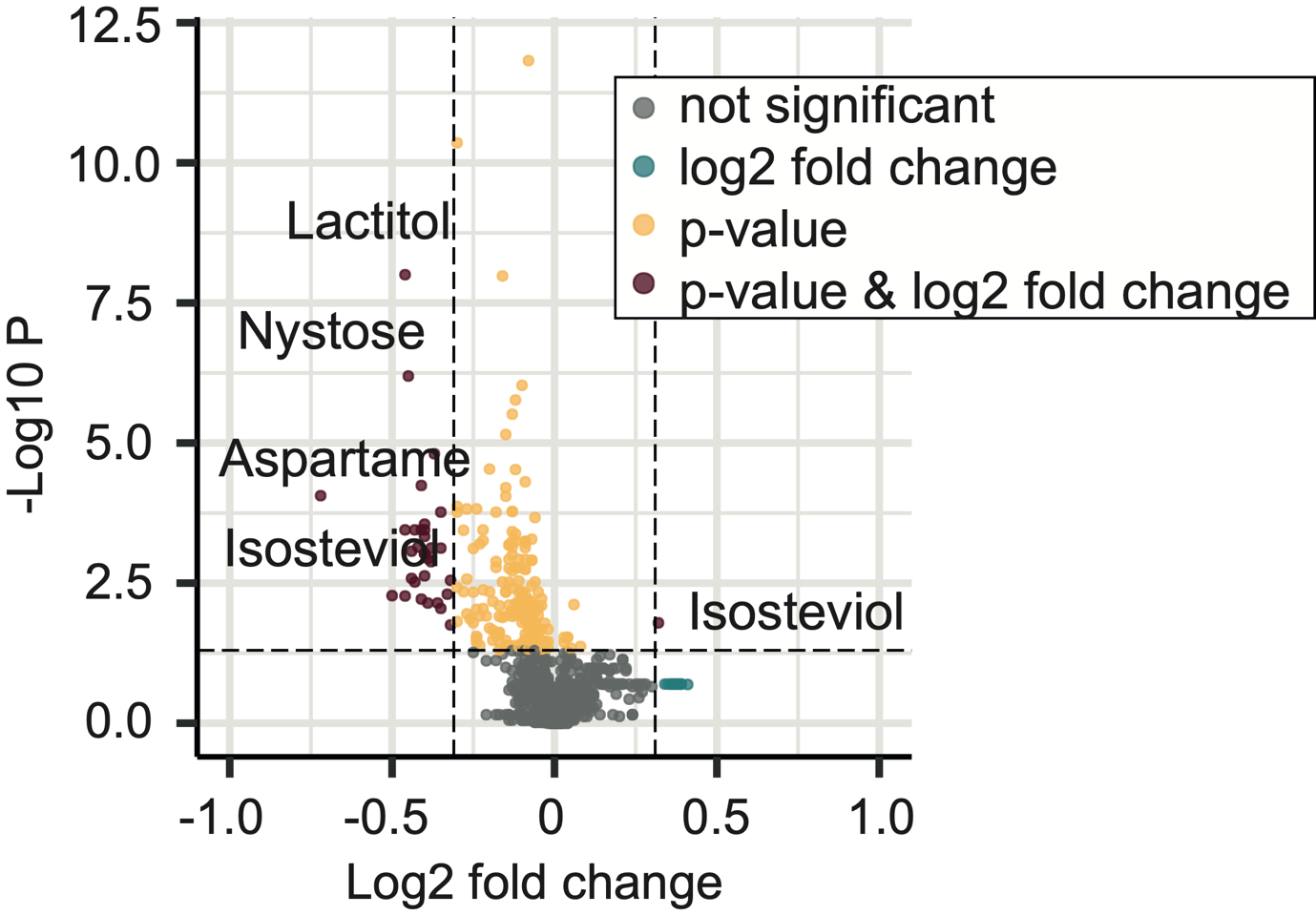

**Supplementary figure 1**. Selection of hits – normalised area under the curve (AUC) across all replicates was pooled and p-value between control and treated wells was computed (Welch's t-test). Median of the log2 fold change of all replicates was computed. Significant hits were determined by setting a threshold of p-value at a confidence interval of 95% and log2 fold change of +/- 0.32 (20% difference in growth)

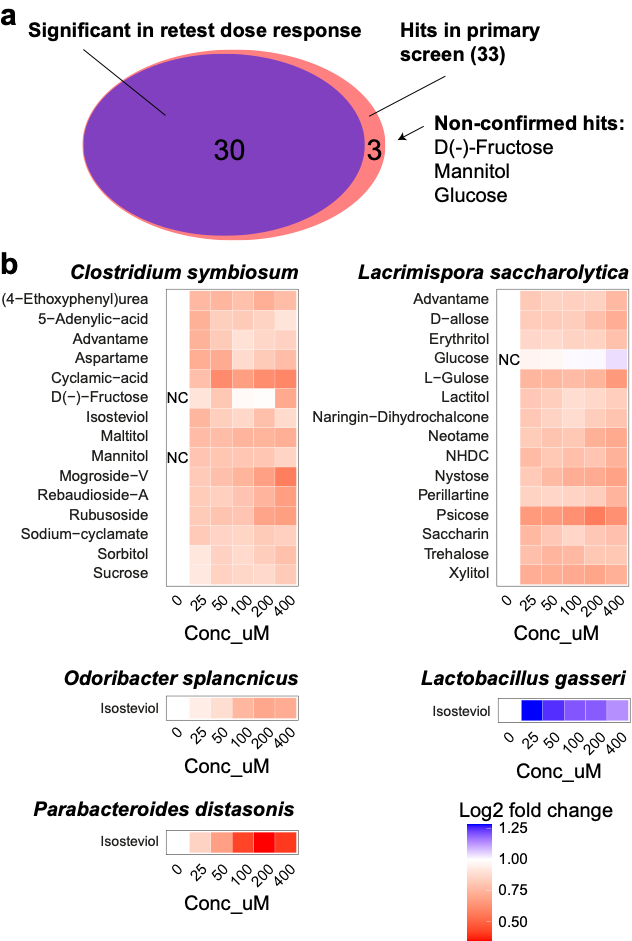

**Supplementary figure 2.** (a) Venn diagram showing the significant hits in the original sweetener-bacteria screen and the retested dose response. (b) Results of the dose response of the retested sweetener screen hits, concentrations tested: 0, 25, 50, 100, 200 and 400 µM. NC = not significant (criteria: min. two conc. significant in p<0.05, N>=3)

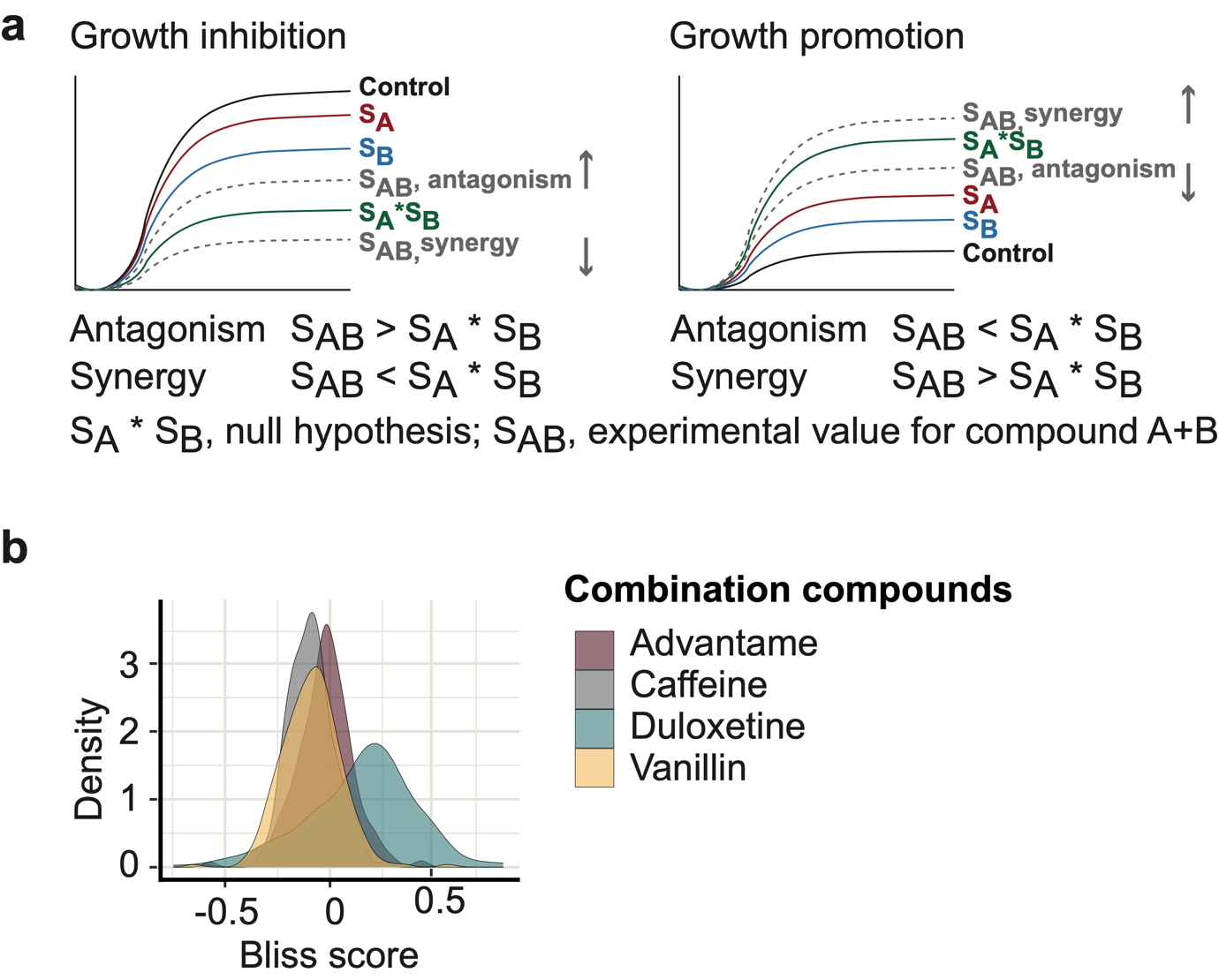

**Supplementary figure 3**. (a) Definition of synergy and antagonism in growth inhibition and growth promotion scenario. (b) Synergistic and antagonistic interactions of sweeteners combined with four commonly used co-occurring compounds.

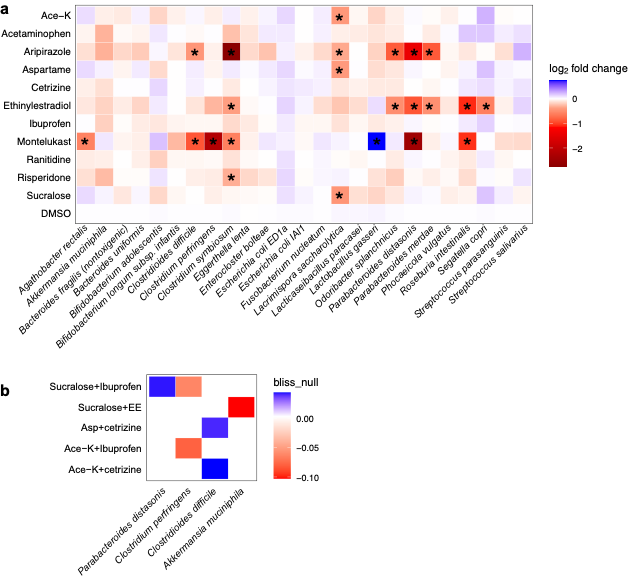

**Supplementary figure 4**. (a) Effect of commonly used drugs and sweeteners on microbial species (* p<0.05) and (b) significant bliss scores of these drugs combined with commonly used sweeteners added to the drug formulation (p<0.05).

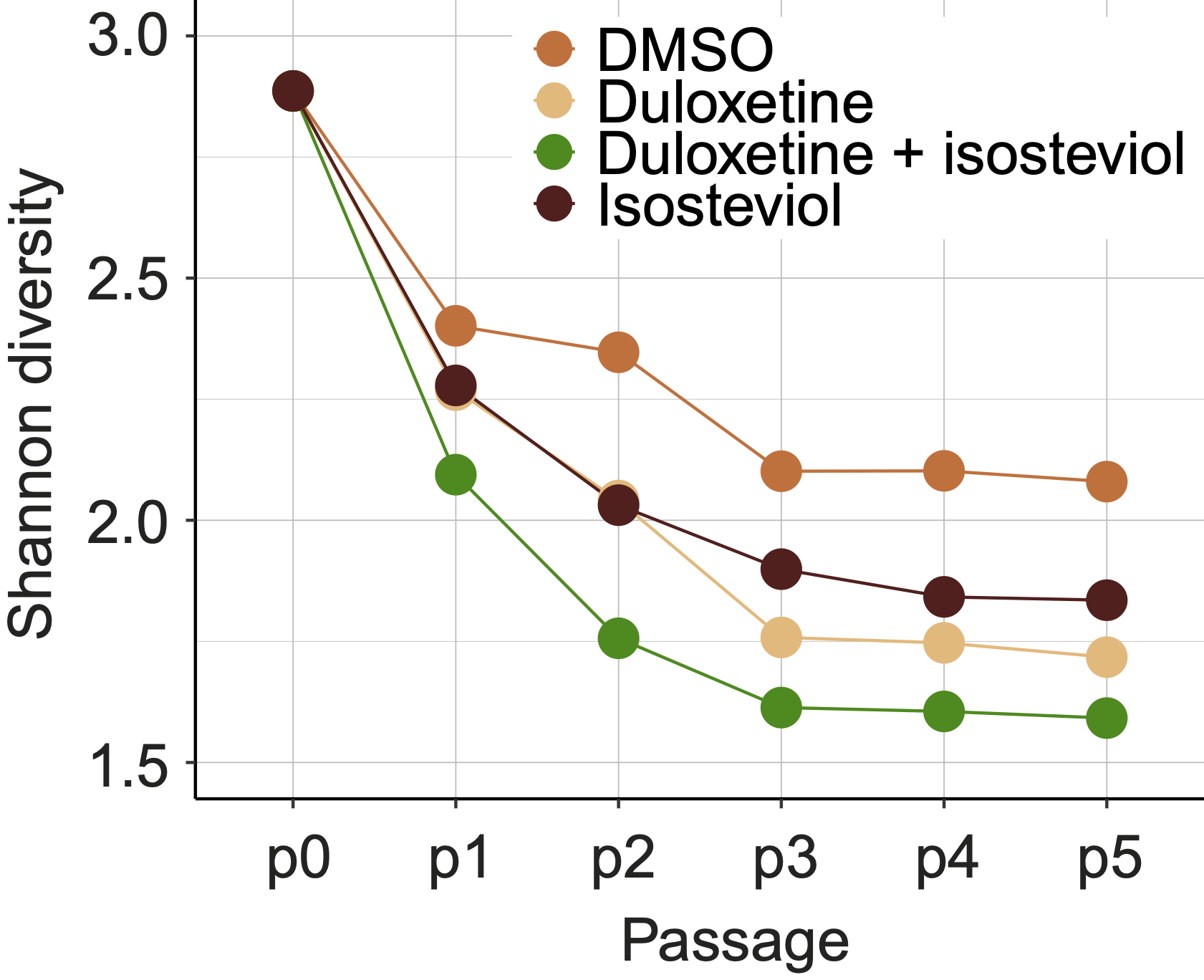

**Supplementary figure 5.** Changes in alpha diversity throughout the passages in compound-treated communities.

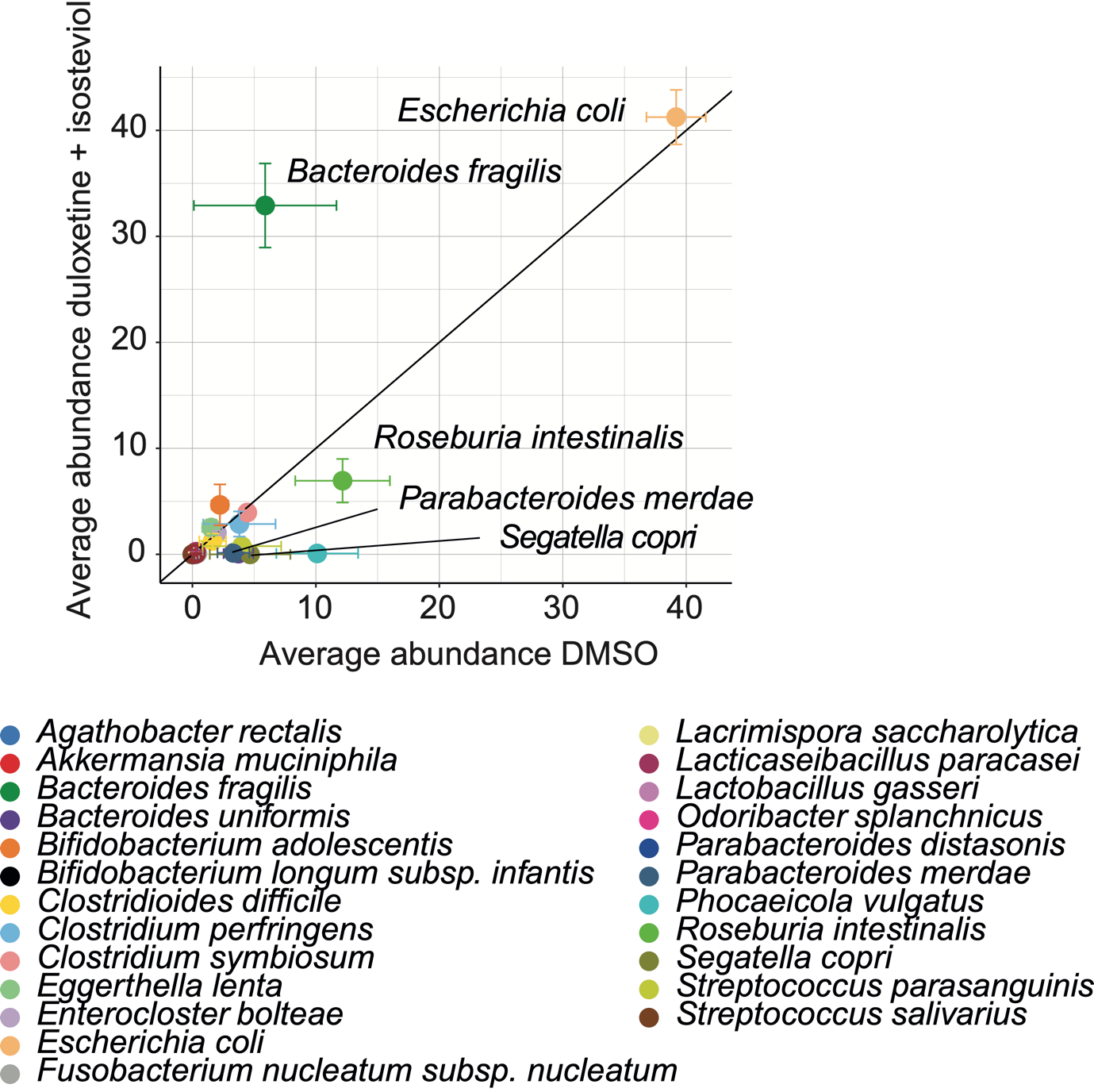

**Supplementary figure 6**. Relative abundance per species in DMSO and isosteviol-duloxetine in fifth (final) passage, species below diagonal are reduced abundance in isosteviol-duloxetine treated communities.

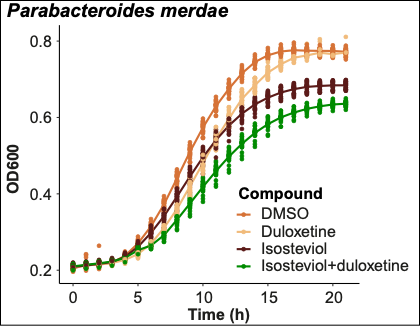

**Supplementary figure 7**. Growth inhibition of *Parabacteroides merdae* by duloxetine, isosteviol and a combination of both as compared to the compound solvent DMSO (1%) at a concentration of 50 µM each. N=24

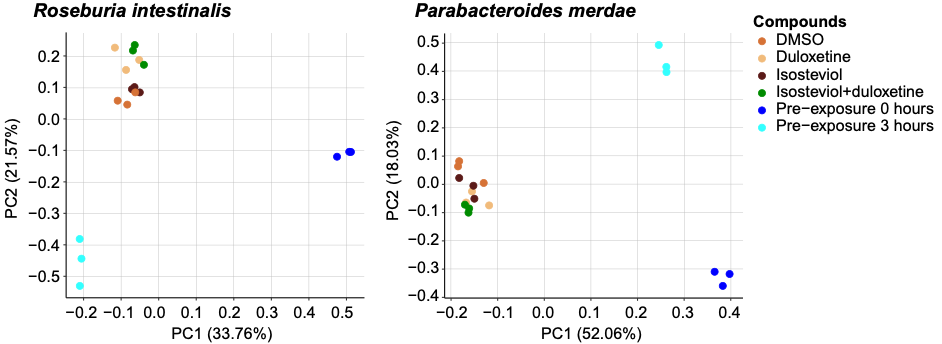

**Supplementary figure 8.** Differences in protein expression in *Roseburia intestinalis* and *Parabacteroides merdae* in different growth stages after culture inoculation. Pre-exposure at 0 hours and at 3 hours without drugs and compounds. Compounds (DMSO, duloxetine, isosteviol and duloxetine + isosteviol, 50 µM each) were added after collection of the 3-hour sample and the bacteria were exposed for 4 hours to each of the compounds. The compound-treated samples were harvested after a total growth time of 7 hours (end of logarithmic growth phase). N=3

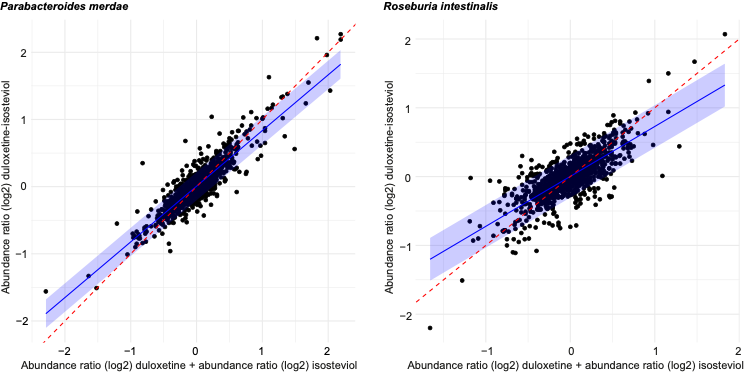

**Supplementary figure 9.** The effect of the combined treatment with duloxetine and isosteviol (y-axis) differs from the addition of the effect of each of the individual compounds in both, *P. merdae* and *R. intestinalis*. This difference is more pronounced in *R. intestinalis* (slope: 0.7257292, R2: 0.6025763) as compared to *P. merdae.* (slope: 0.828918, R2: 0.8049975).

a
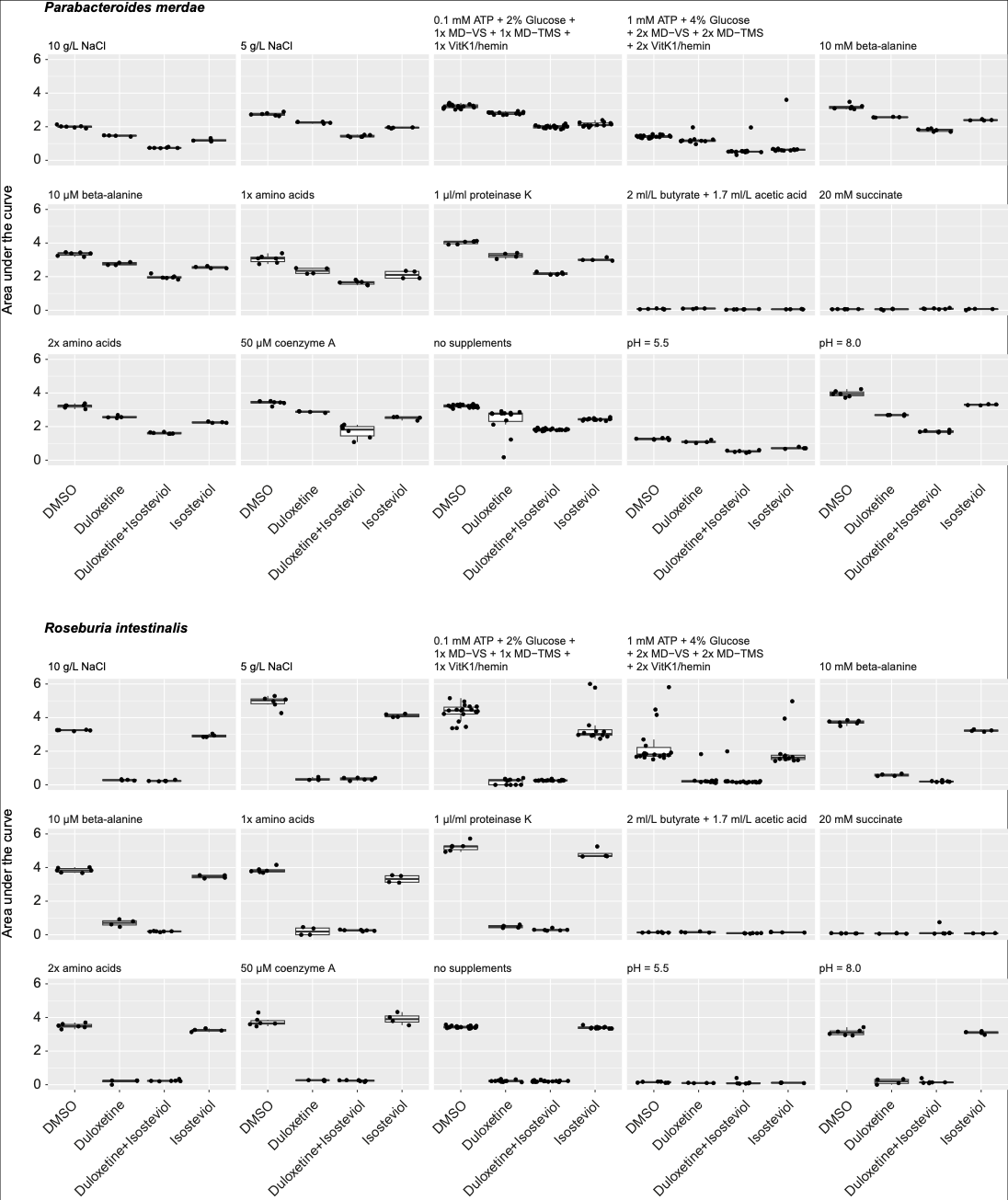

b
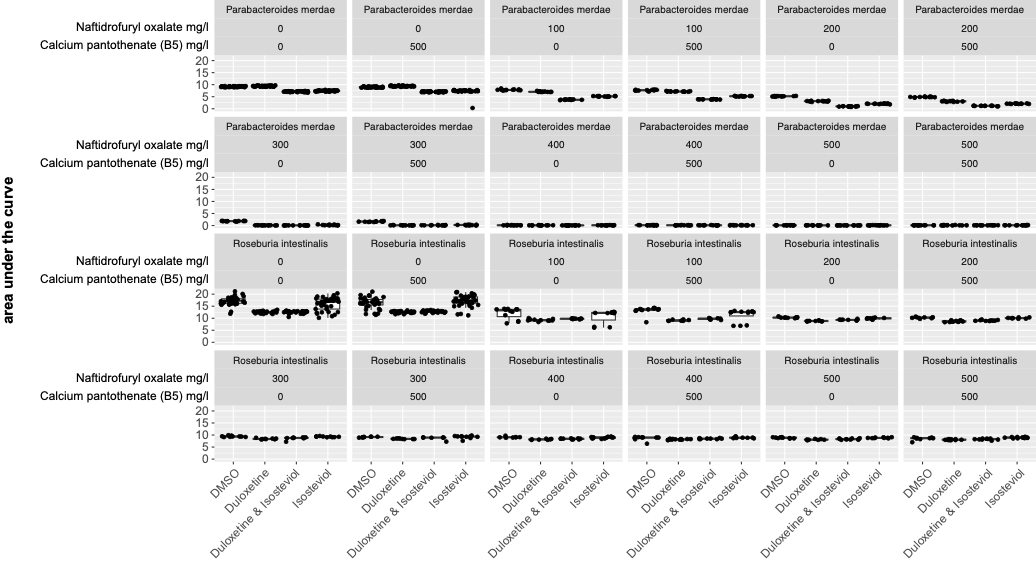

**Supplementary figure 10.** (a) A selection of the supplements was tested for their effect on the synergy between isosteviol and duloxetine. Although several supplements were able to improve or inhibit the growth of *R. intestinalis* and *P. merdae*, neither of them alone nor in combination was able to specifically affect the synergy. N = 6. (b) Different concentrations of naftidrofuryl (nafronyl) oxalate (an inhibitor of pantothenate synthetase, panC) and calcium pantothenate were tested in different combinations in the two bacteria exposed to isosteviol, duloxetine and the combination, N=12. Naftidrofuryl oxalate inhibited the growth, while calcium pantothenate was unable to restore it.

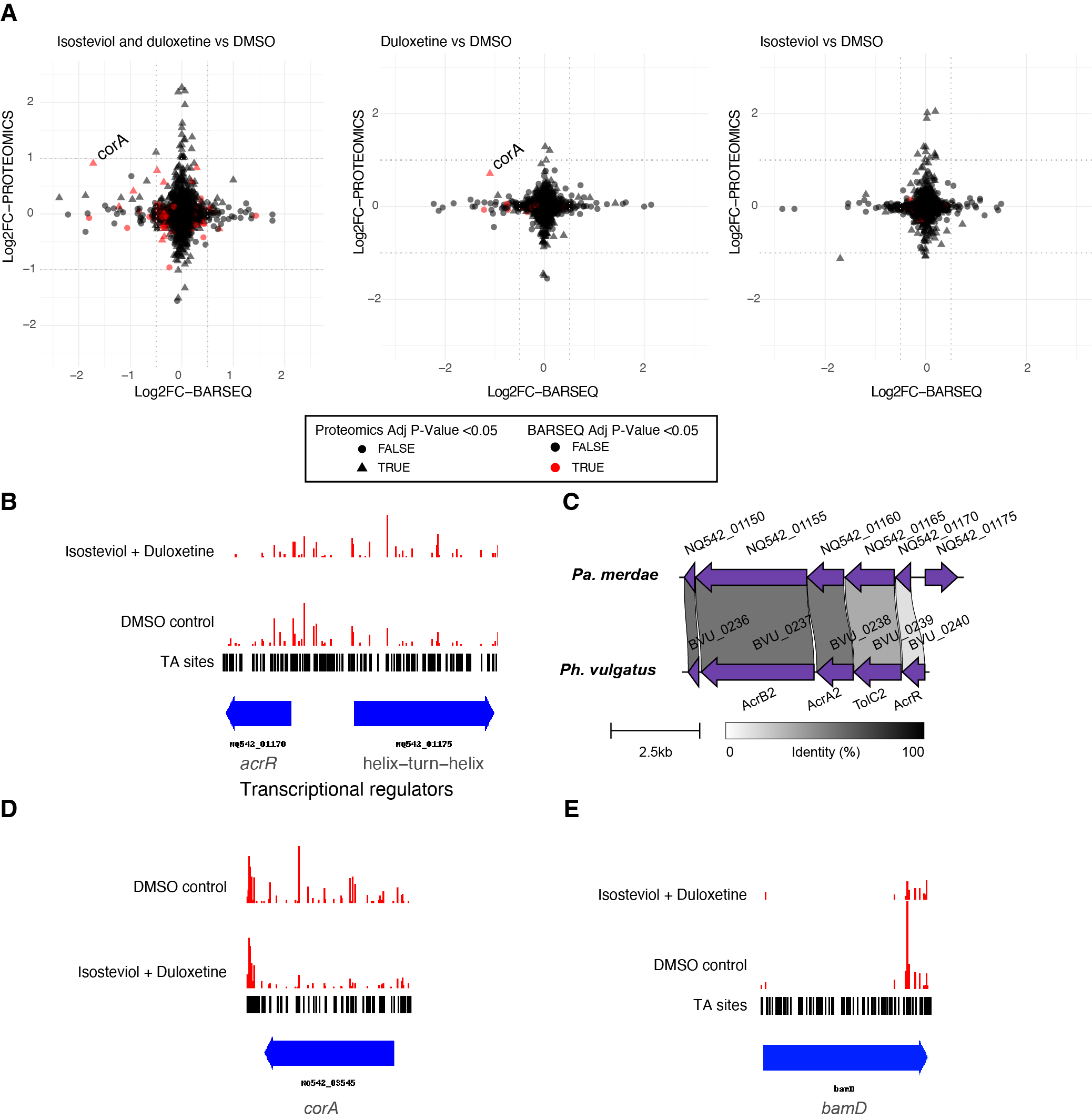

**Supplementary figure 11. A.** Comparison of gene fitness (BarSeq) with the expression values of the corresponding proteins (proteomics) under duloxetine, isosteviol and co-exposure compared to DMSO control. Only genes detected and matched in both datasets are shown (as per Supporting Table X). Each dot represents a gene, with significant hits in BarSeq indicated in red, and significant in proteomic hits as a triangle. CorA magnesium transporter stands as an upregulated gene with strong negative fitness. **B.** Representative map of insertions in *acrR* transporter gene (negative fitness), a divergent helix-turn-helix regulator gene (positive fitness) and intergenic region (negative fitness). The blue arrows represent genes, and the reads from transposon insertions are represented in red. The presence of TA sites (preferential transposon insertion site) is shown as a black track. **C.** Representative schematic of the genetic neighbourhood of the transcriptional regulator gene *acrR* located proximal to *acrB-tolC* genes, as its orthologue recently characterised as modulator of efflux transport in *B. vulgatus*^1^ made with clinker^2^. This suggests that regulation of efflux is involved in detoxification of duloxetine and combination. **D.** Representative map of insertions in *corA* **E.** Representative map of insertions at the C-terminal domain of the outer membrane protein assembly factor BamD. Although a cut-off of insertions in central 80% of the coding gene was applied when computing gene fitness score, several *bamD* partial knock outs show a decreased fitness under duloxetine, and isosteviol + duloxetine.

1.	Hibberd, M. C. *et al.* The effects of micronutrient deficiencies on bacterial species from the human gut microbiota. *Sci Transl Med* **9**, (2017).

2.	Gilchrist, C. L. M. & Chooi, Y. H. Clinker & clustermap.js: Automatic generation of gene cluster comparison figures. *Bioinformatics* **37**, 2473–2475 (2021).

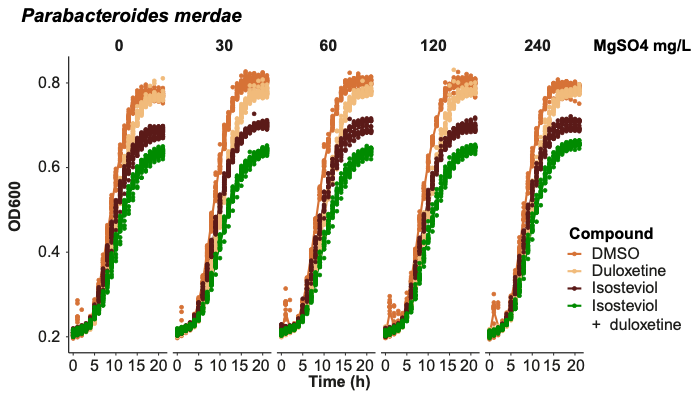

**Supplementary figure 12.** Supplementation of *P. merade* with magnesium sulfate (MgSO4) at different concentrations. MgSO4 did not alter growth up to a concentration of 240 mg/L in mGAM medium. N=24.

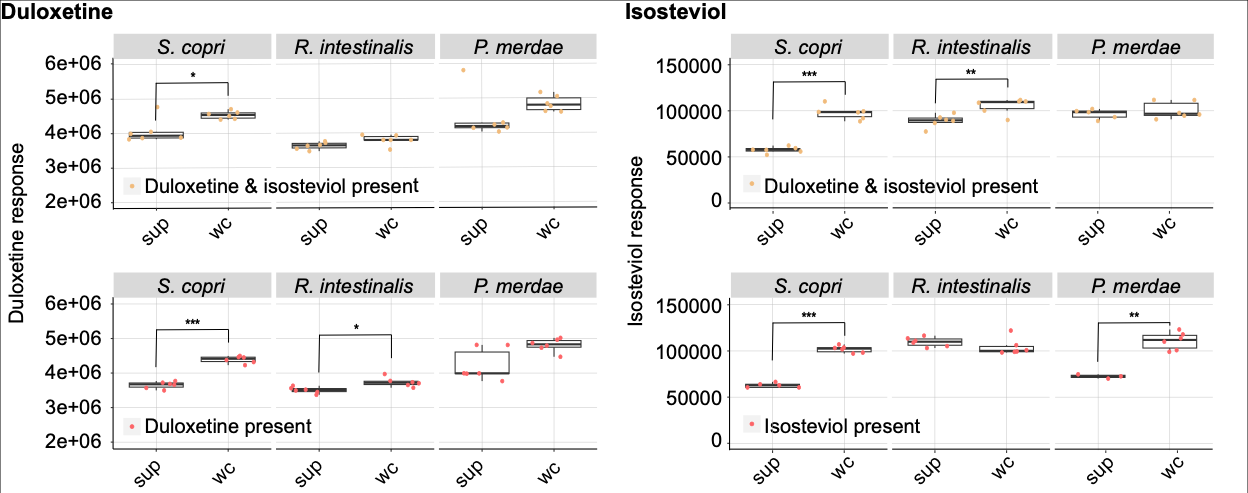

**Supplementary figure 13.** Bioaccumulation of duloxetine and isosteviol in *Segatella copri*, *R. intestinalis* and *P. merdae* when applied alone and in combination. *S. copri* bioaccumulates each, duloxetine and isosteviol, equally when applied alone and in combination. *R. intestinalis* bioaccumulates duloxetine only in absence of isosteviol and isosteviol only in presence of duloxetine. However, the quantitative difference is small. *P. merdae* bioaccumulates isosteviol only in absence of duloxetine. * p<0.05, ** p<0.01, *** p<0.001, N=6, wc=whole culture, sup=supernatant.

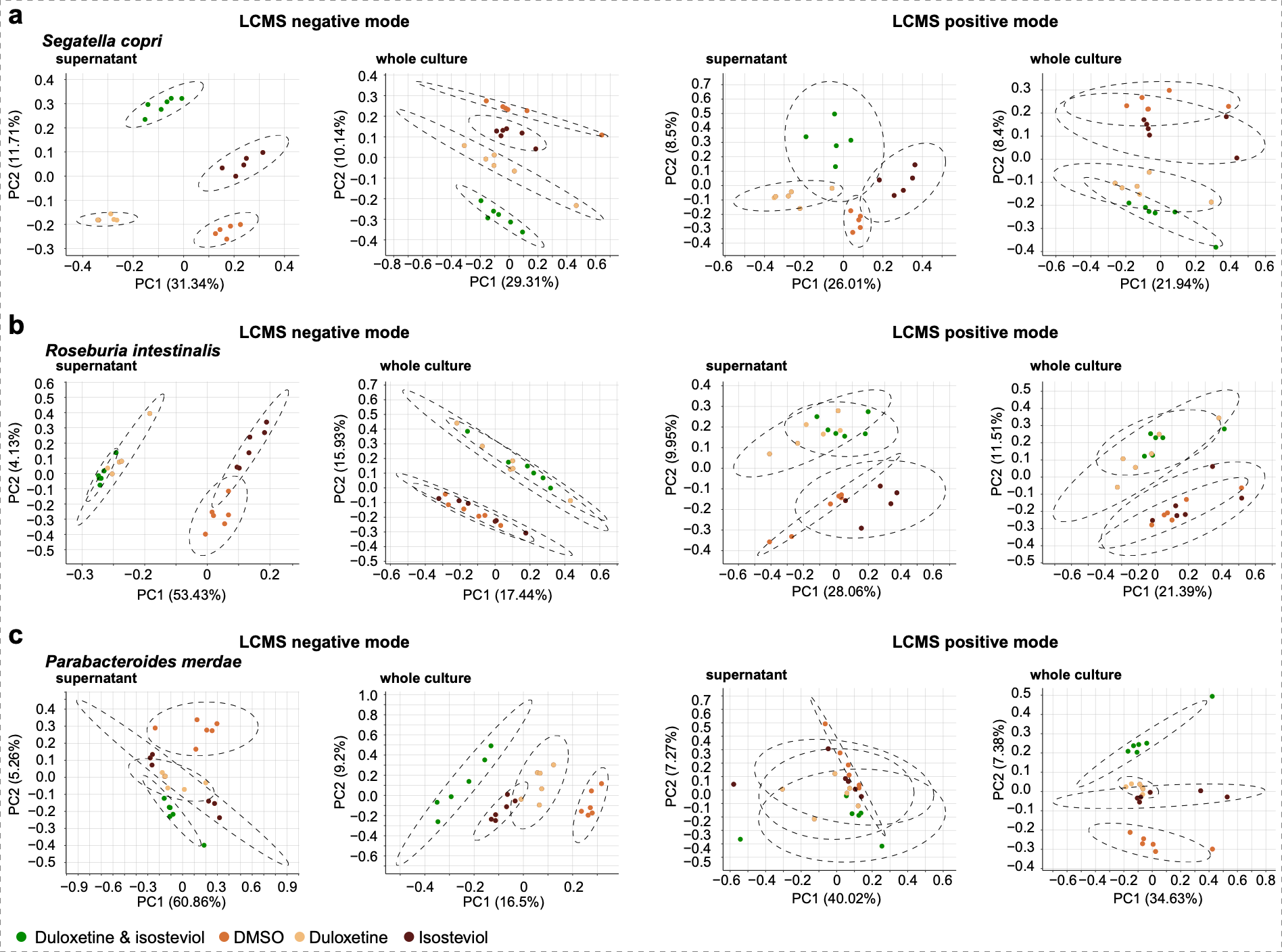

**Supplementary figure 14.** Altered metabolite production and secretion in *S. copri*, *R. intestinalis* and *P. merdae* upon overnight exposure to duloxetine, isosteviol and the combination of both (50 µM each in mGAM) as compared to the solvent (DMSO). N=6.

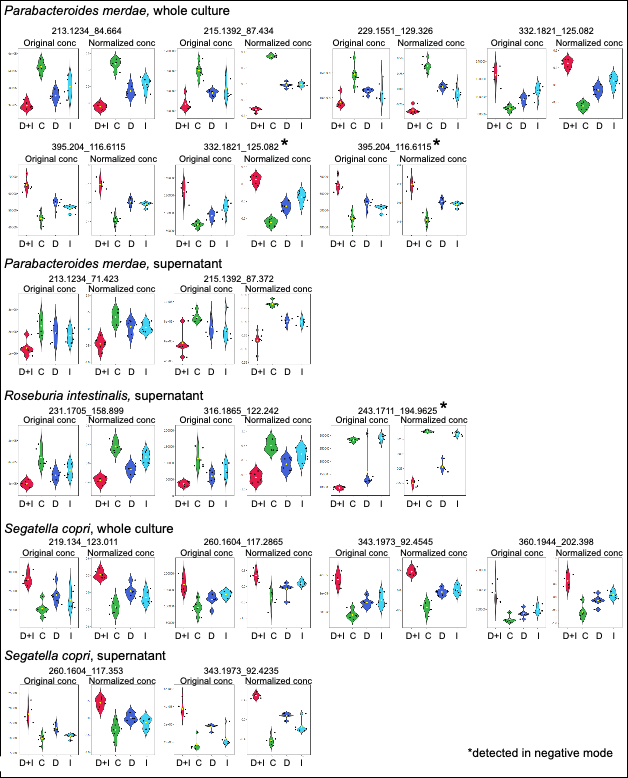

**Supplementary figure 15.** Metabolites affected from combined exposure to duloxetine and isosteviol of *R. intestinalis*, *P merdae* and *S. copri*. D+I = Duloxetine + isosteviol, C = control, D = duloxetine, I = isosteviol, N=6.

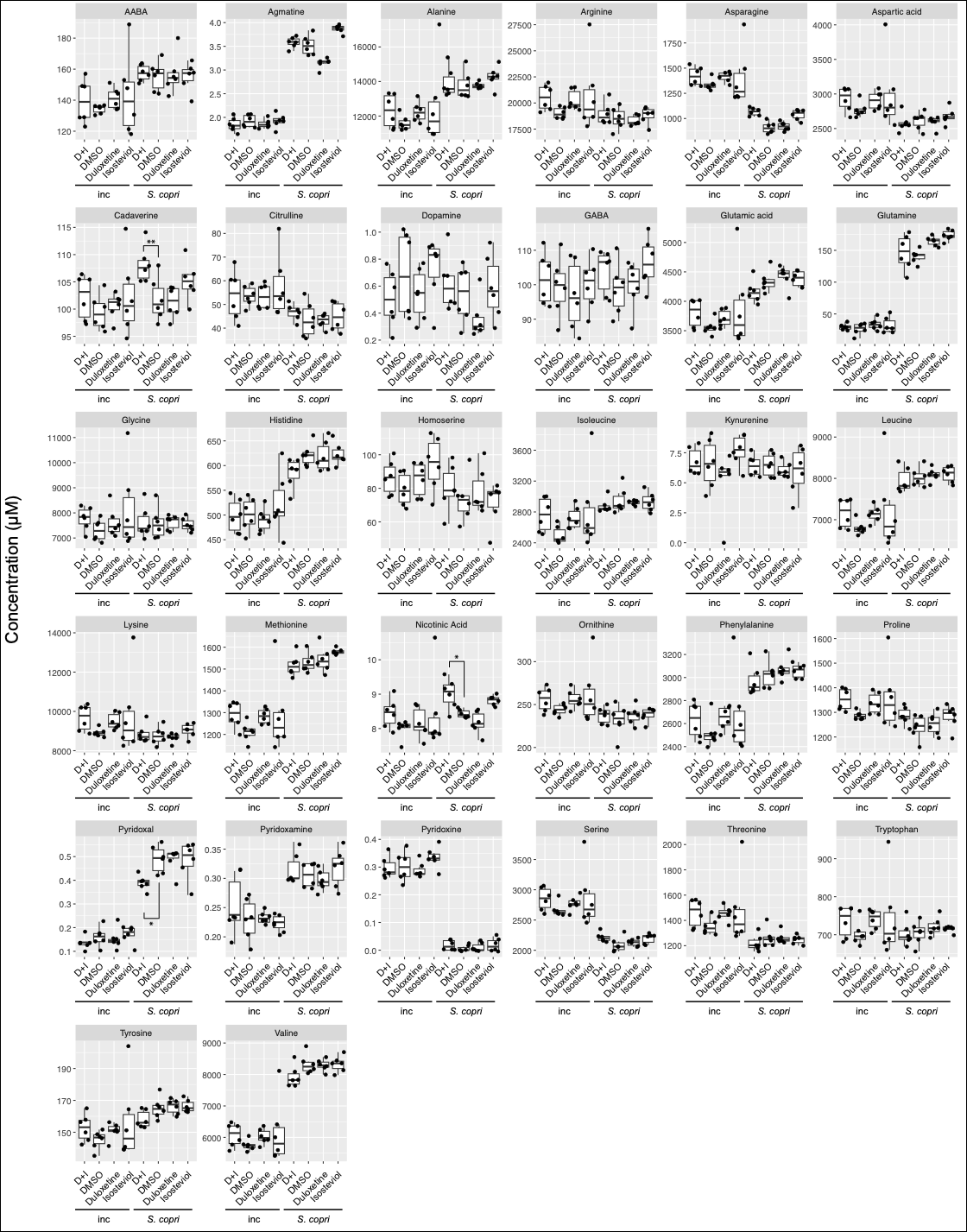

**Supplementary figure 16.** Changes in the concentrations of 32 amino acids and derivates in *Segatella copri* upon overnight exposure to duloxetine, isosteviol and the combination of both (50 µM each). D+I = Duloxetine + isosteviol, inc = incubation control (mGAM medium plus compounds, without bacteria). N=6, * = p<0.05.

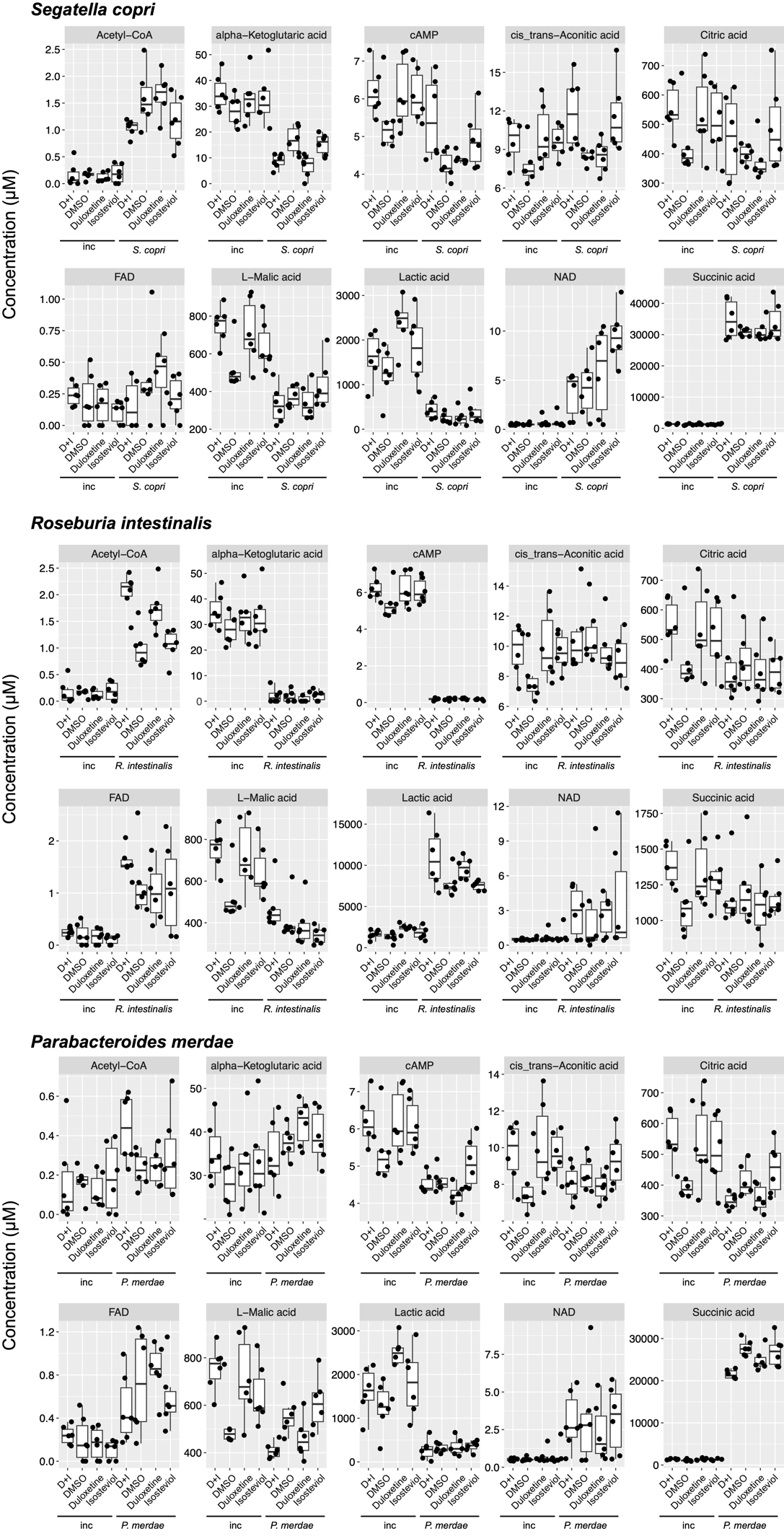

**Supplementary figure 17.** Changes in the concentrations of organic acids in *Segatella copri*, *Roseburia intestinalis* and *Parabacteroides merdae* upon overnight exposure to duloxetine, isosteviol and the combination of both (50 µM each). D+I = Duloxetine + isosteviol, inc = incubation control (mGAM medium plus compounds, without bacteria). N=6, * = p<0.05, ** = p<0.01, *** = p<0.001, **** = p<0.0001.

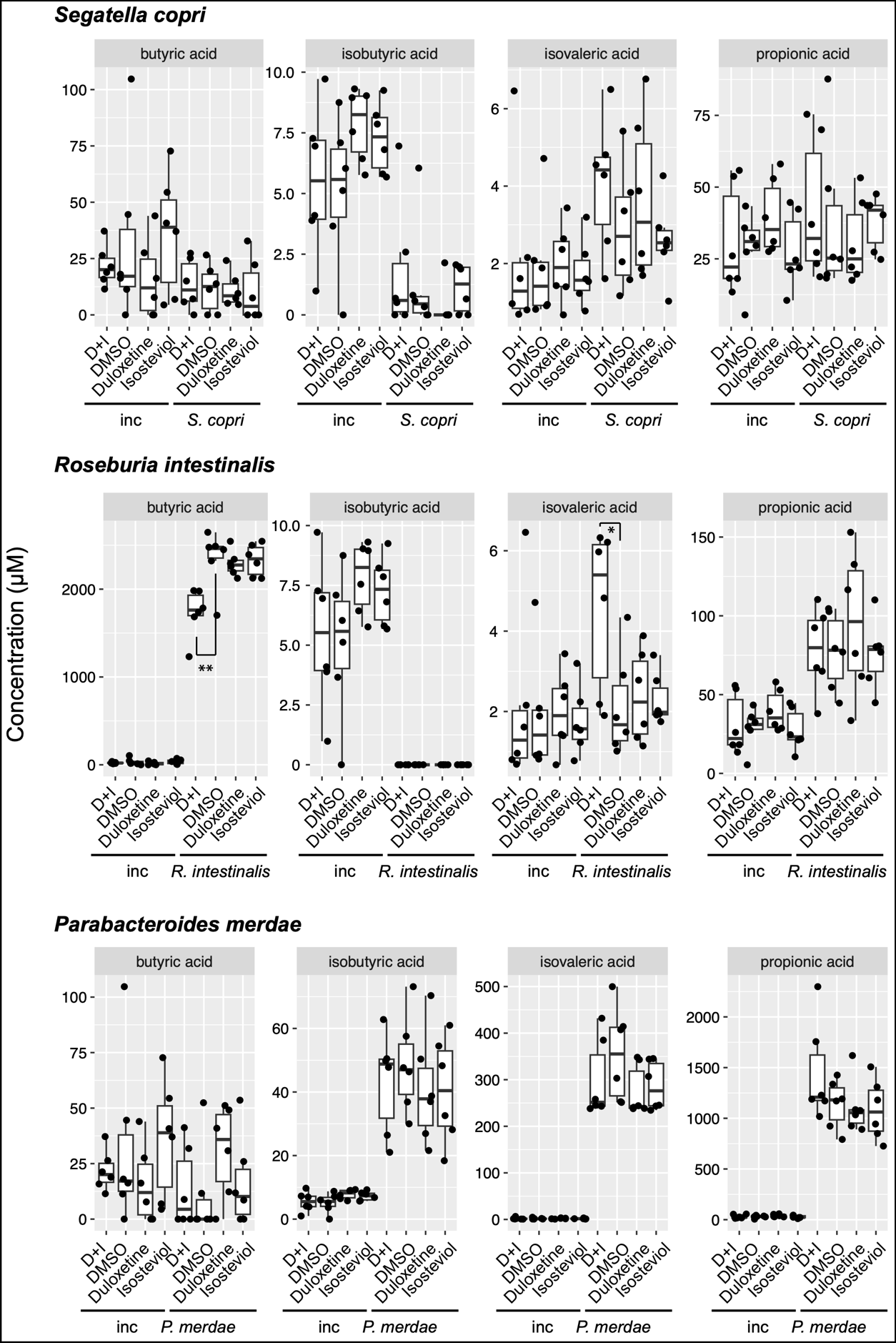

**Supplementary figure 18.** Changes in the concentrations of short chain fatty acids in *Segatella copri*, *Roseburia intestinalis* and *Parabacteroides merdae* upon overnight exposure to duloxetine, isosteviol and the combination of both (50 µM each). D+I = Duloxetine + isosteviol, inc = incubation control (mGAM medium plus compounds, without bacteria). N=6, * = p<0.05, ** = p<0.01, *** = p<0.001, **** = p<0.0001.

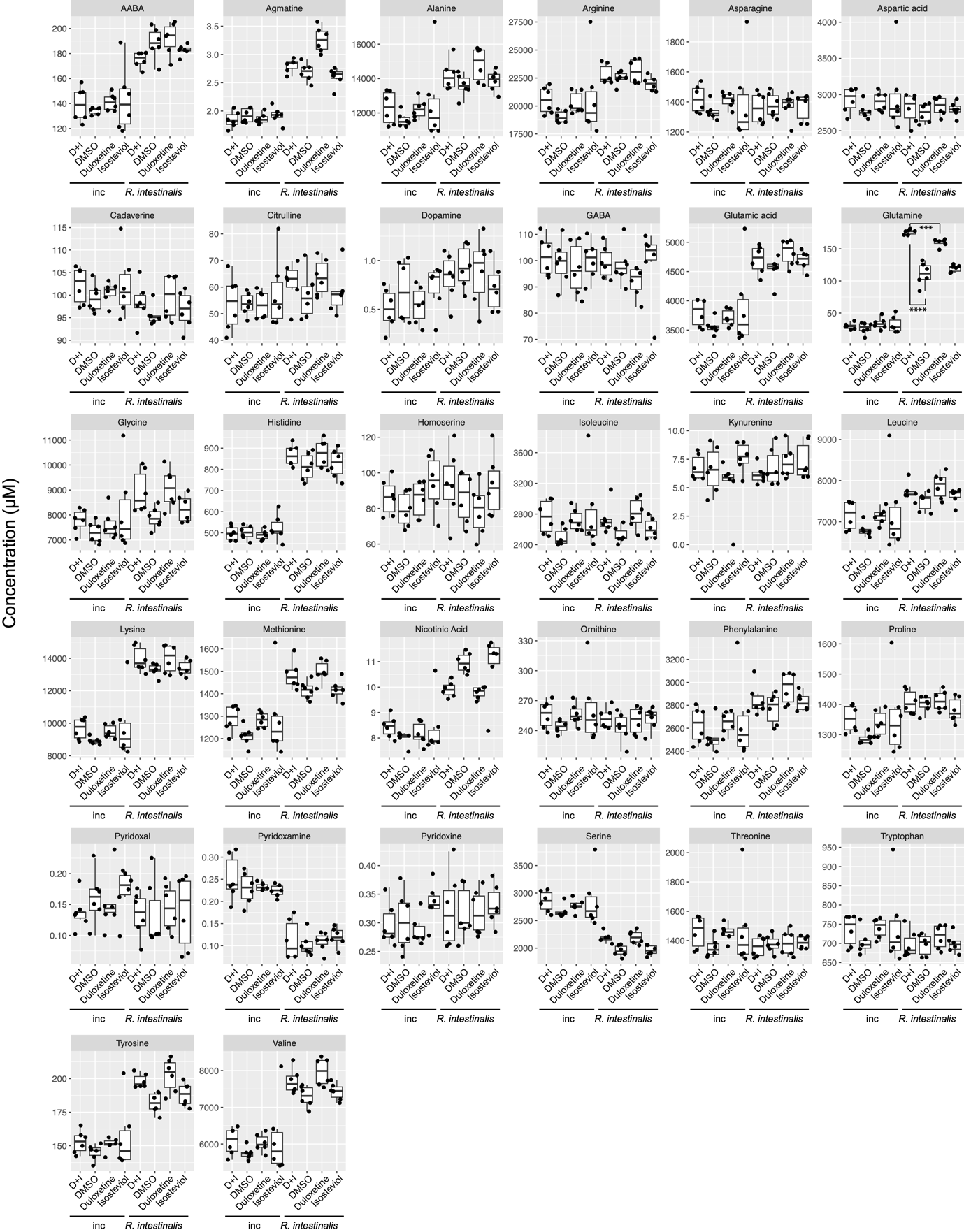

**Supplementary figure 19.** Changes in the concentrations of 32 amino acids and derivates in *Roseburia intestinalis* upon overnight exposure to duloxetine, isosteviol and the combination of both (50 µM each). D+I = Duloxetine + isosteviol, inc = incubation control (mGAM medium plus compounds, without bacteria). N=6, * = p<0.05, ** = p<0.01, *** = p<0.001, **** = p<0.0001.

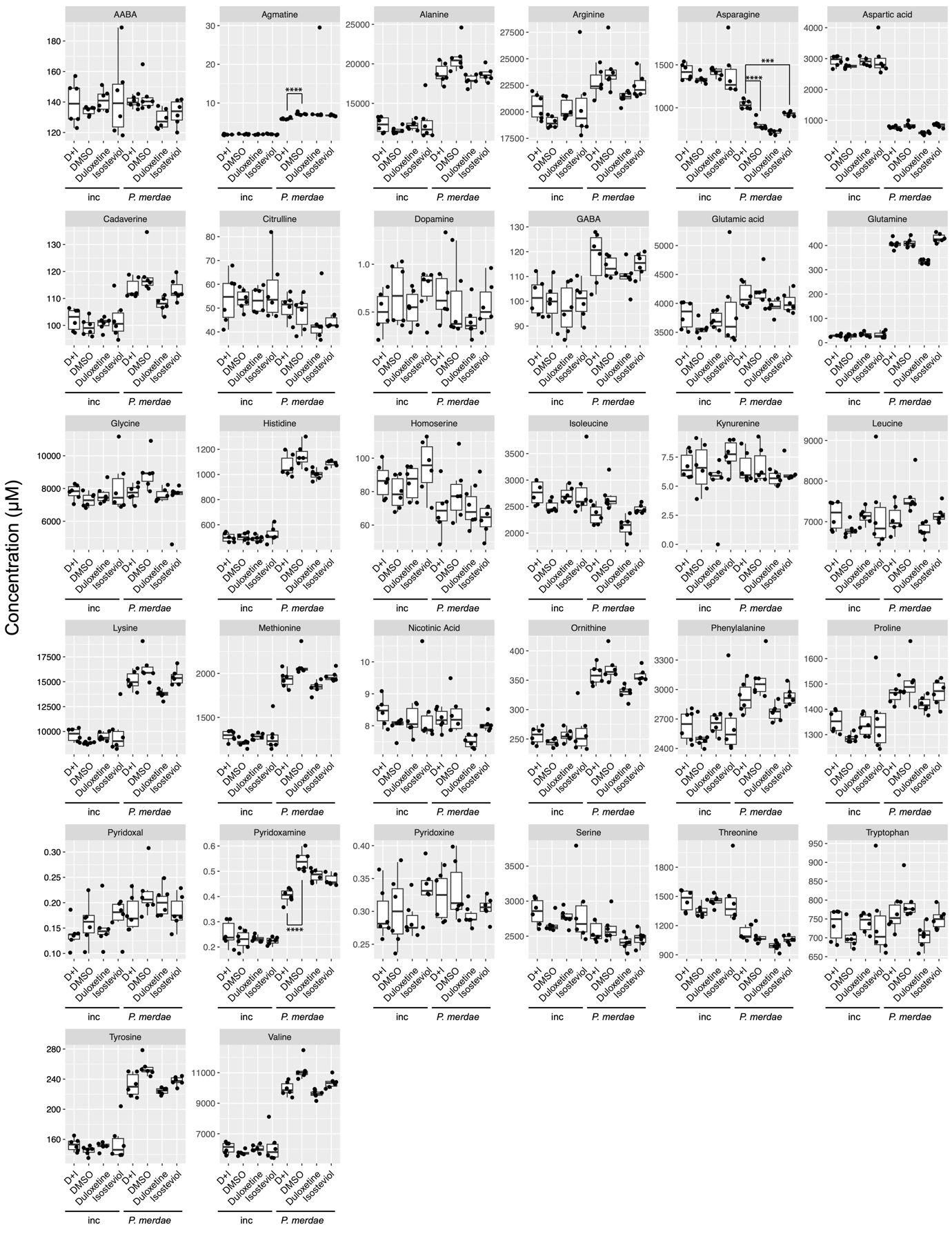

**Supplementary figure 20.** Changes in the concentrations of 32 amino acids and derivates in *Parabacteroides merdae* upon overnight exposure to duloxetine, isosteviol and the combination of both (50 µM each). D+I = Duloxetine + isosteviol, inc = incubation control (mGAM medium plus compounds, without bacteria). N=6, * = p<0.05, ** = p<0.01, *** = p<0.001, **** = p<0.0001.

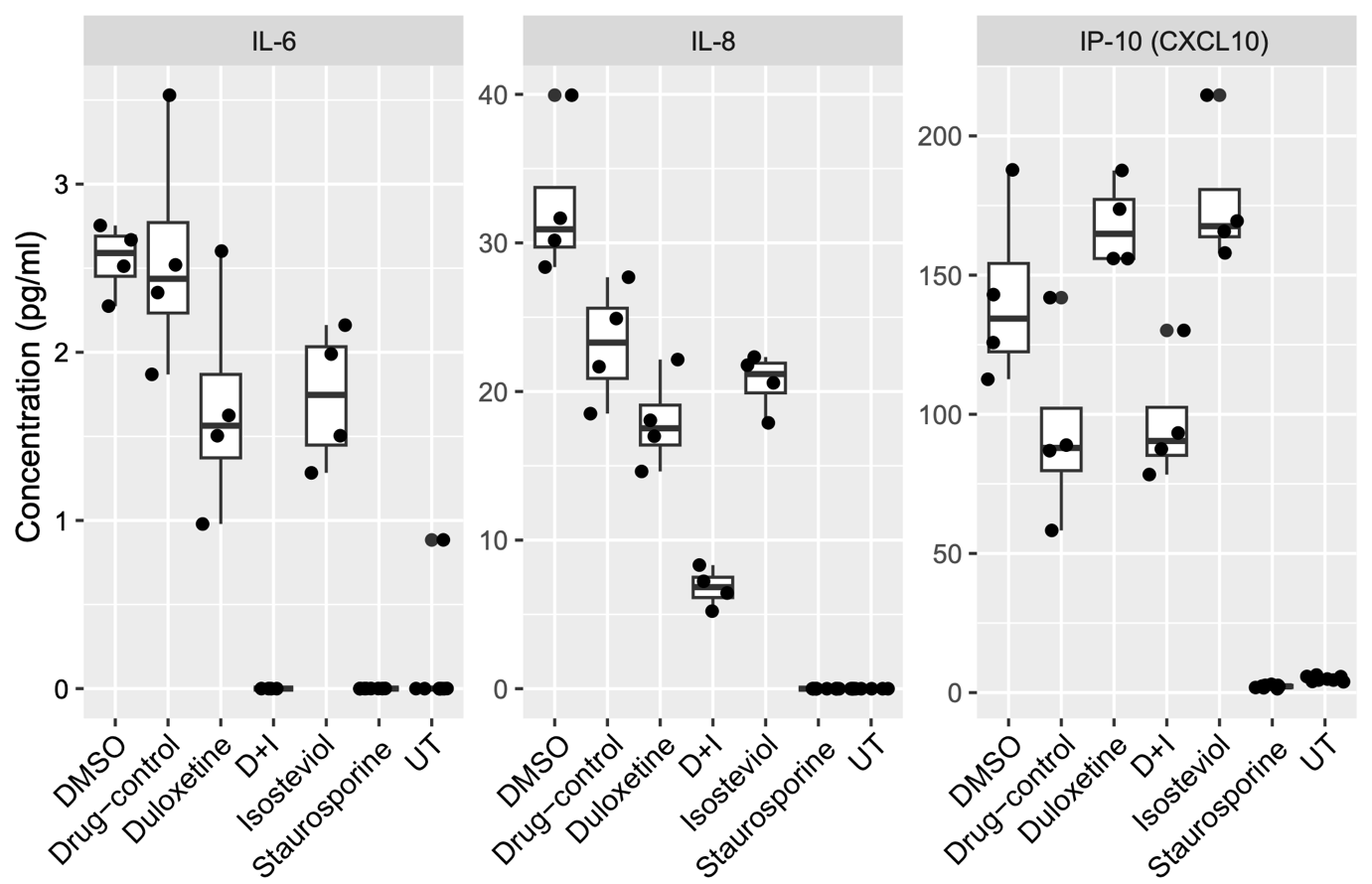

**Supplementary figure 21.** Secretion of the cytokines interleukin-6 (IL-6), interleukin-8 (IL-8) and the interferon-γ inducible protein 10 kDa (CXCL10) by Caco-2 cells as a response to contact with spent medium. IL-6 and IL-8 secretion is significantly reduced for the supernatant of duloxetine and isosteviol co-treated communities, as compared to the DMSO and drug control. The 25- bacteria community was treated with duloxetine, isosteviol, and duloxetine + isosteviol (D+I). Controls are DMSO (solvent, 0.2%) treated community and the supernatant of the solvent-treated community with 50 µM duloxetine and isosteviol, added after harvesting (drug control). Staurosporine and DMEM alone (UT) does not induce any of the cytokines. N>6.

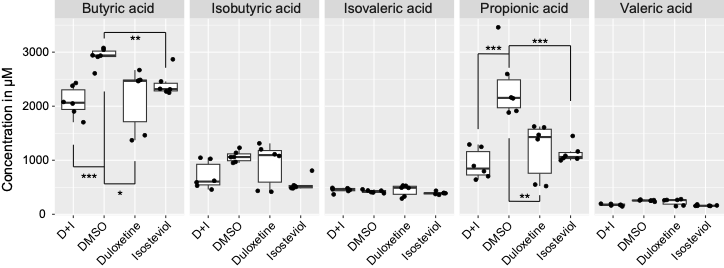

**Supplementary figure 22.** Concentration of the five short chain fatty acids in the supernatant of the 25 bacteria community. The supernatant contains ~3 mM butyric acid and 2 mM propionic acid. Both SCFAs are significantly decreased in duloxetine, isosteviol and duloxetine plus isosteviol (D+I) treated community supernatants. N=6, * = p<0.05, ** = p<0.01, *** = p<0.001.

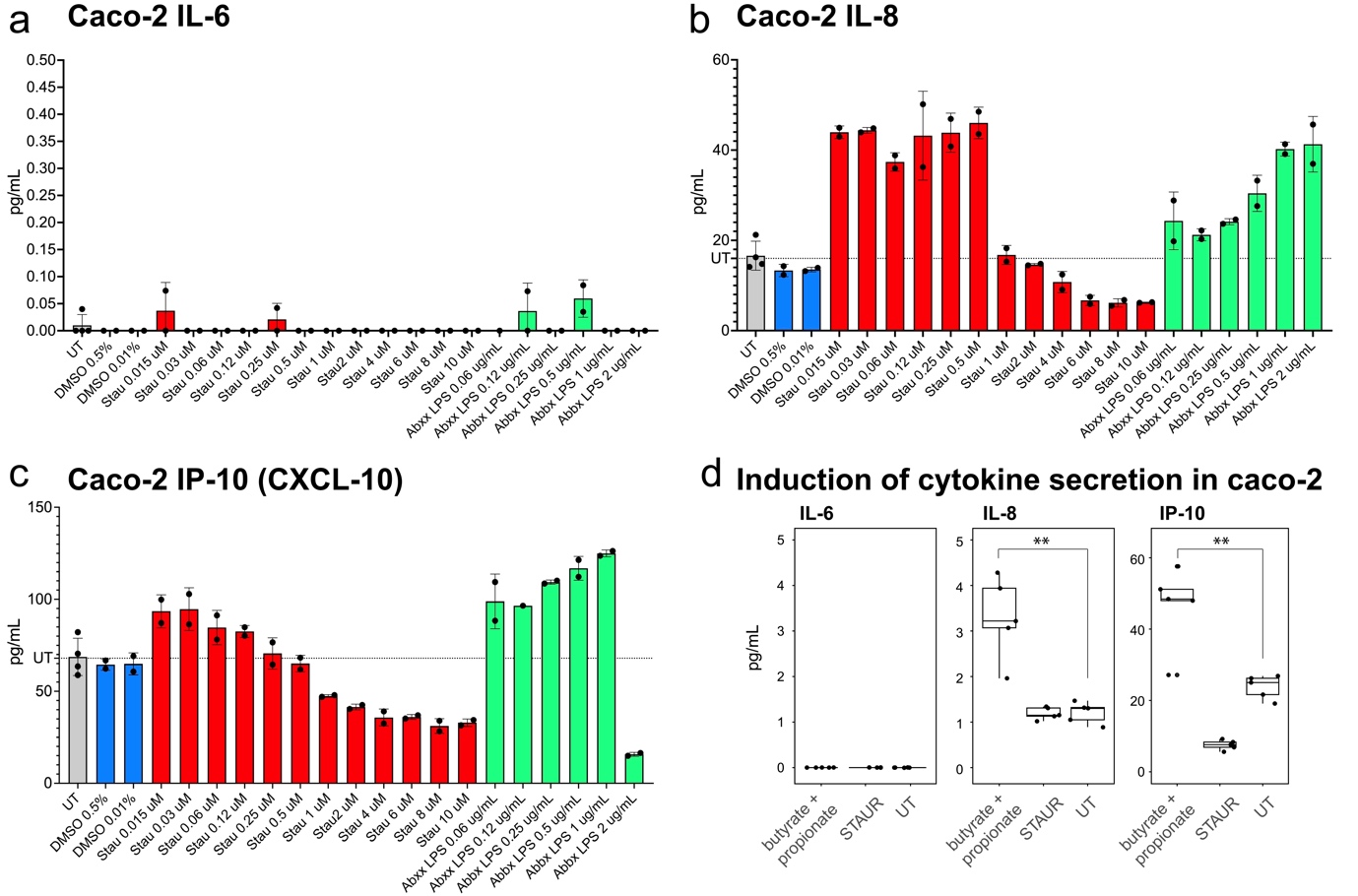

**Supplementary figure 23**. (a-c) Secretion of IL-6, IL-8 and IP-10 after exposure to different concentrations of staurosporine and LPS in Caco-2 cells. Secretion of each cytokine is below that of untreated cells (UT) for staurosporine (stau) concentrations above 1 µM. A concentration of 5 µM or more is required for apoptosis induction. (d) Induction of cytokine secretion in caco-2 cells after treatment with 3 mM butyrate + 2 mM propionic acid, compared to 5 µM staurosporine (STAUR) and untreated cells. ** p<0.01, N=5

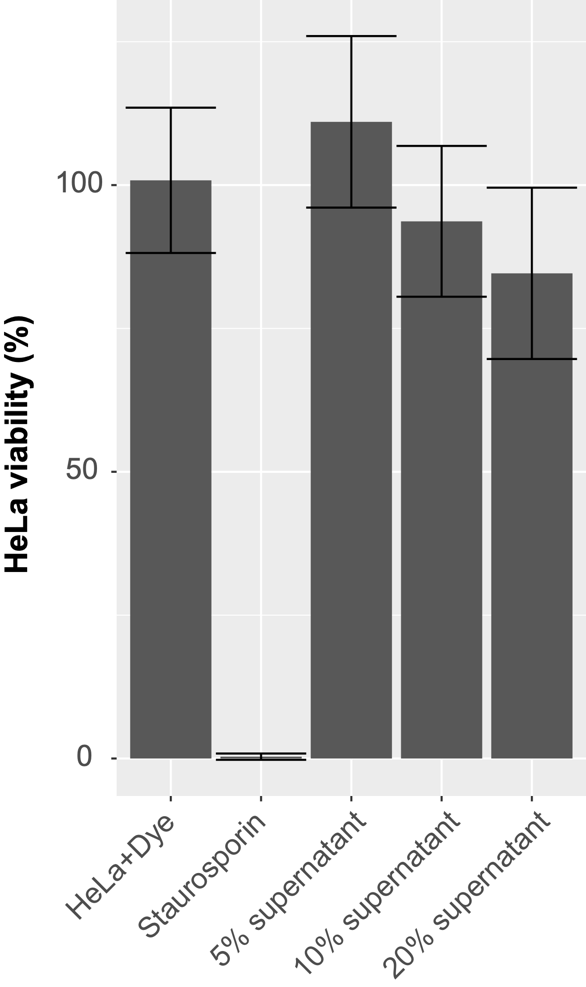

**Supplementary figure 24.** Viability test for HeLa cells exposed to 5, 10 and 20 % spent media from the 25 species community, done in triplicates.

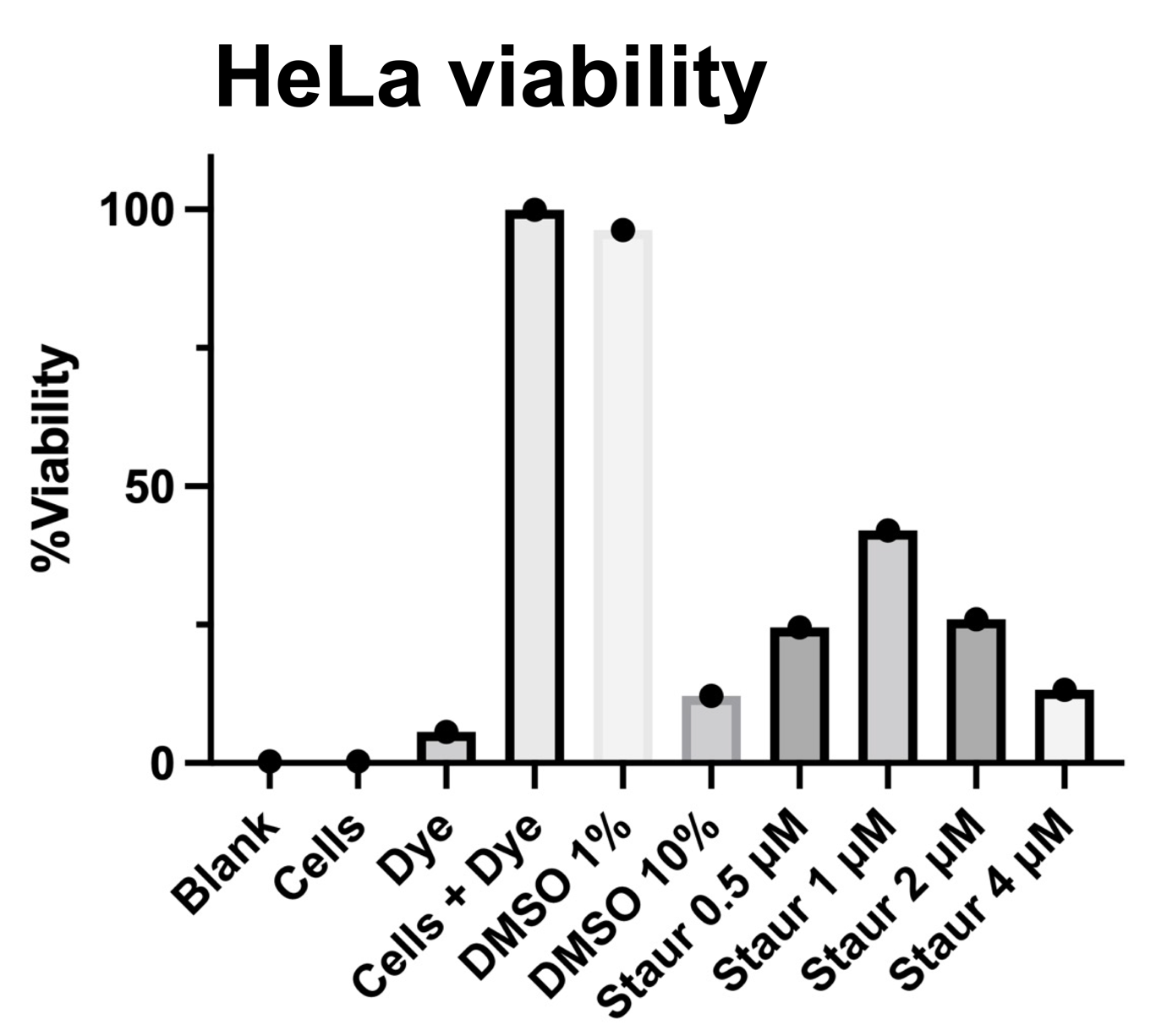

**Supplementary figure 25**. HeLa cell viability after treatment with different concentrations of staurosporine for apoptosis induction.

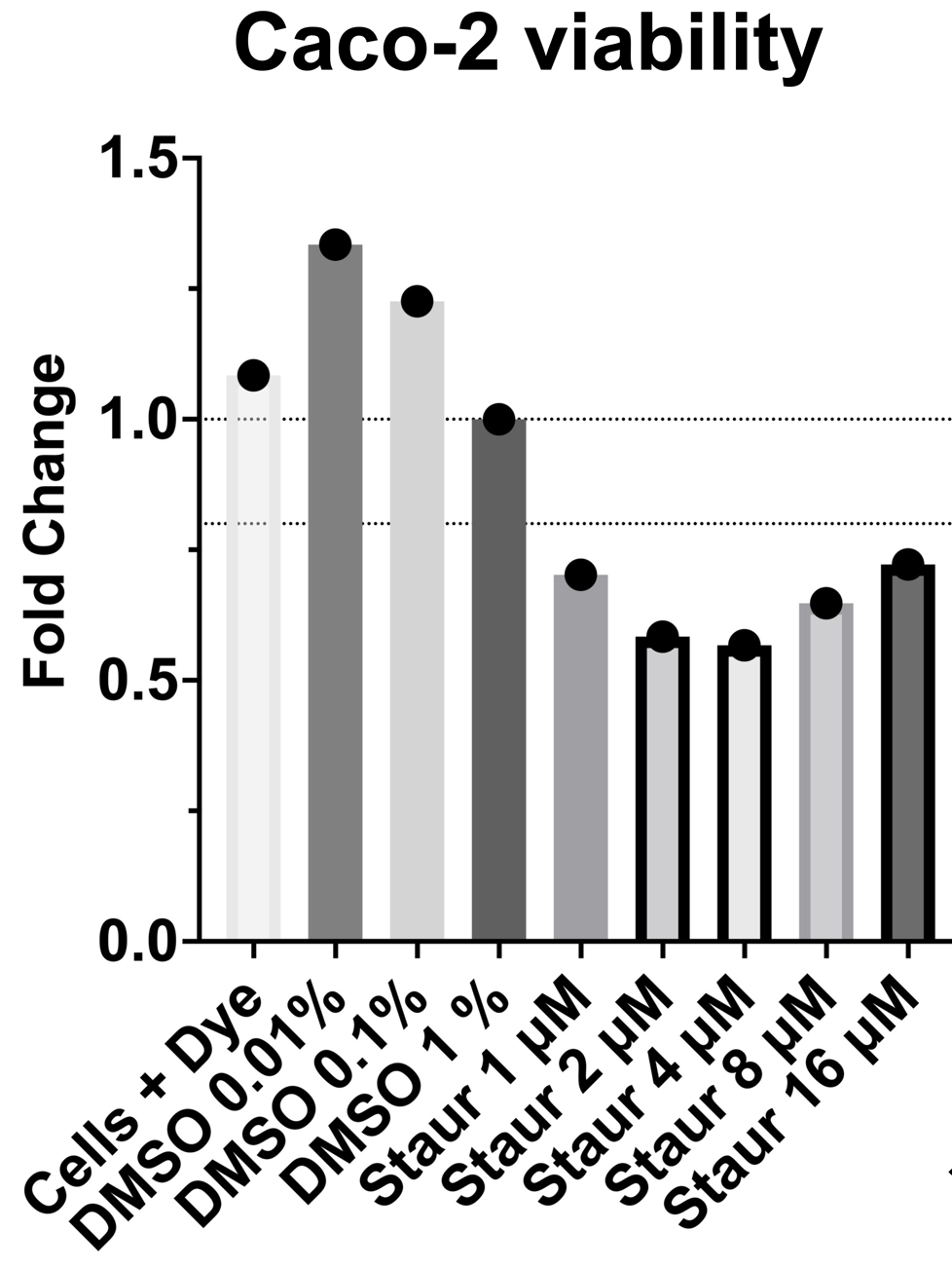

**Supplementary figure 26**. Caco-2 viability after treatment with different concentrations of staurosporine for apoptosis induction.
